# TMEM164 is an acyltransferase that forms ferroptotic polyunsaturated ether phospholipids

**DOI:** 10.1101/2022.07.06.498872

**Authors:** Alex Reed, Timothy Ware, Haoxin Li, J. Fernando Bazan, Benjamin F. Cravatt

**Affiliations:** Department of Chemistry, The Scripps Research Institute, San Diego, CA 92037; ℏ bioconsulting, llc, Stillwater, MN 55082, and %Unit for Structural Biology, VIB-UGent Center for Inflammation Research, Technologiepark 71, 9052 Ghent, Belgium

## Abstract

Ferroptosis is an iron-dependent form of cell death driven by the oxidation of polyunsaturated (PUFA) phospholipids. Large-scale genetic screens have pointed to a specialized role for PUFA ether phospholipids (ePLs) in promoting ferroptosis. Our understanding of the enzymes involved in PUFA ePL production, however, remains incomplete. Here we show using a combination of pathway mining of genetic dependency maps, AlphaFold-guided structure predictions, and targeted lipidomics that the uncharacterized transmembrane protein TMEM164 – genetic ablation of which has been shown to protect cells from ferroptosis – is a cysteine active-center enzyme that selectively transfers C20:4 acyl chains from phosphatidylcholine to lyso-ePLs to furnish PUFA-ePLs. TMEM164-null cells show substantial reductions in PUFA-ePLs, but not PUFA ester phospholipids, supporting that the selective suppression of PUFA-ePLs is sufficient to protect cells from ferroptosis and designating TMEM164 as a key enzyme specifically responsible for regulating this class of lipids.

## Introduction

Polyunsaturated phospholipids (PUFA-PLs) serve critical functions in cell biology that include modulation of the structure (e.g., fluidity) of membrane bilayers and as precursors for an array of signaling molecules (e.g., eicosanoids, endocannabinoids)^1^. PUFA-PLs are susceptible to enzymatic and non-enzymatic oxidation and controlling the extent of PUFA-PL oxidation is critical for cell homeostasis and viability^2^. For instance, excess PUFA-PL oxidation can lead to ferroptosis, an iron-dependent form of non-apoptotic cell death^3^. A key enzyme involved in counteracting excessive PUFA oxidation is glutathione peroxidase 4 (GPX4), which converts lipid hydroperoxides to lipid alcohols, and the genetic or pharmacological disruption of GPX4 promotes ferroptosis in susceptible cell types^4,5^. Ferroptosis has been implicated in a wide range of human diseases, including cancer and various degenerative disorders^6^. Accordingly, mapping the biochemical pathways that regulate ferroptosis may not only improve our understanding of the mechanistic underpinnings for this cell death process, but also offer therapeutic targets for enhancing or suppressing ferroptosis sensitivity in specific disease settings.

Recent genome-wide CRISPR–Cas9 suppressor screens have revealed a central role for PUFA ether-PLs in regulating ferroptosis, specifically identifying genes encoding peroxisomal biogenesis proteins or peroxisomal enzymes involved in ePL biosynthesis as pro-ferroptotic^7,8^. Mechanistic studies further revealed that cancer cells can convert to a ferroptosis-resistant state *in vivo* that is associated with reductions in PUFA-ePL content^7^. Concurrent with this work, independent investigations have begun to showcase the potential for genome-wide mapping of co-essential genes to identify orphan members of biochemical pathways, including the discovery that the previously uncharacterized membrane protein TMEM189 plays a key role in ePL biosynthesis through acting as a plasmanylethanolamine desaturase^9^. Additional studies support this function^10,11^, as well as the potential for TMEM189, at least in certain settings, to regulate ferroptotic activity^8^.

Despite the specialized role emerging for PUFA-ePLs in ferroptosis, it is notable that the pro-ferroptotic ePL enzymes identified so far appear to exhibit generalized substrate scopes that lack acyl chain specificity, which contrasts with other lipid metabolic enzymes implicated in ferroptosis such as ACSL4 and LPCAT3 (or MBOAT5) that display a strong preference for PUFA (C20:4 or arachidonate) substrates^12–14^. Here, we show using a combination of pathway reconstruction based on genetic dependency maps, structural predictions by AlphaFold, lipidomics, and biochemical and cell biological assays that the uncharacterized transmembrane protein TMEM164 acts as a cysteine-dependent, C20:4-preferring acyltransferase that generates PUFA-ePLs in human cells. Notably, TMEM164 uses C20:4-PC (rather than C20:4-CoA) as an acyl chain donor, providing molecular annotation to an enzymatic activity first described over 30+ years ago in the literature^15–18^. We further show that genetic disruption of TMEM164 leads to substantial and selective reductions in C20:4-ePLs, with a concomitant increase in saturated and monounsaturated ePLs, providing a mechanistic explanation for the protection from ferroptosis observed in TMEM164-null cells. These data, taken together, underscore the importance of PUFA-ePLs in ferroptosis and designate TMEM164 as a key enzyme specifically responsible for regulating this class of lipids.

## Results

### TMEM164 has co-dependency relationships with both ePL and PUFA lipid pathways

We hypothesized that additional genes involved in PUFA-ePL metabolism might show codependency relationships with established ePL enzymes in the Cancer Dependency Map^9,19^. Among such genes, TMEM164, which codes for an uncharacterized integral membrane protein, caught our attention because it displayed a co-dependency relationship with not only ePL enzymes (e.g., AGPS, FAR1, TMEM189), but also with key enzymes specifically involved in PUFA (C20:4) lipid metabolism (e.g., ACSL4, LPCAT3) (**Fig. 1a**). GO enrichment analysis of the top 40 co-dependencies for TMEM164 also supported a specific association with ether lipid metabolism (**Fig. 1b**). Interestingly, TMEM164 further showed a stronger connectivity to ACSL4 and LPCAT3 compared to most of the other established ePL enzymes (e.g., FAR1, AGPS, GNPAT) (**Fig. 1c**), suggesting that TMEM164 may represent a biochemical link between ePL and PUFA lipid pathways (**Fig. 1d**). Also consistent with this hypothesis, recent genome-wide CRISPR–Cas9 suppressor screens have revealed that genetic disruption of TMEM164, like other ePL and PUFA lipid pathway members, impairs ferroptosis^7,20,21^. We independently verified that TMEM164-null (sgTMEM164) 786-O human renal cell carcinoma cells show greatly reduced sensitivity to ferroptosis induction by the GPX4 inhibitors ML210 and RSL3 (**Extended Data Fig. 1**).

**Figure 1.**
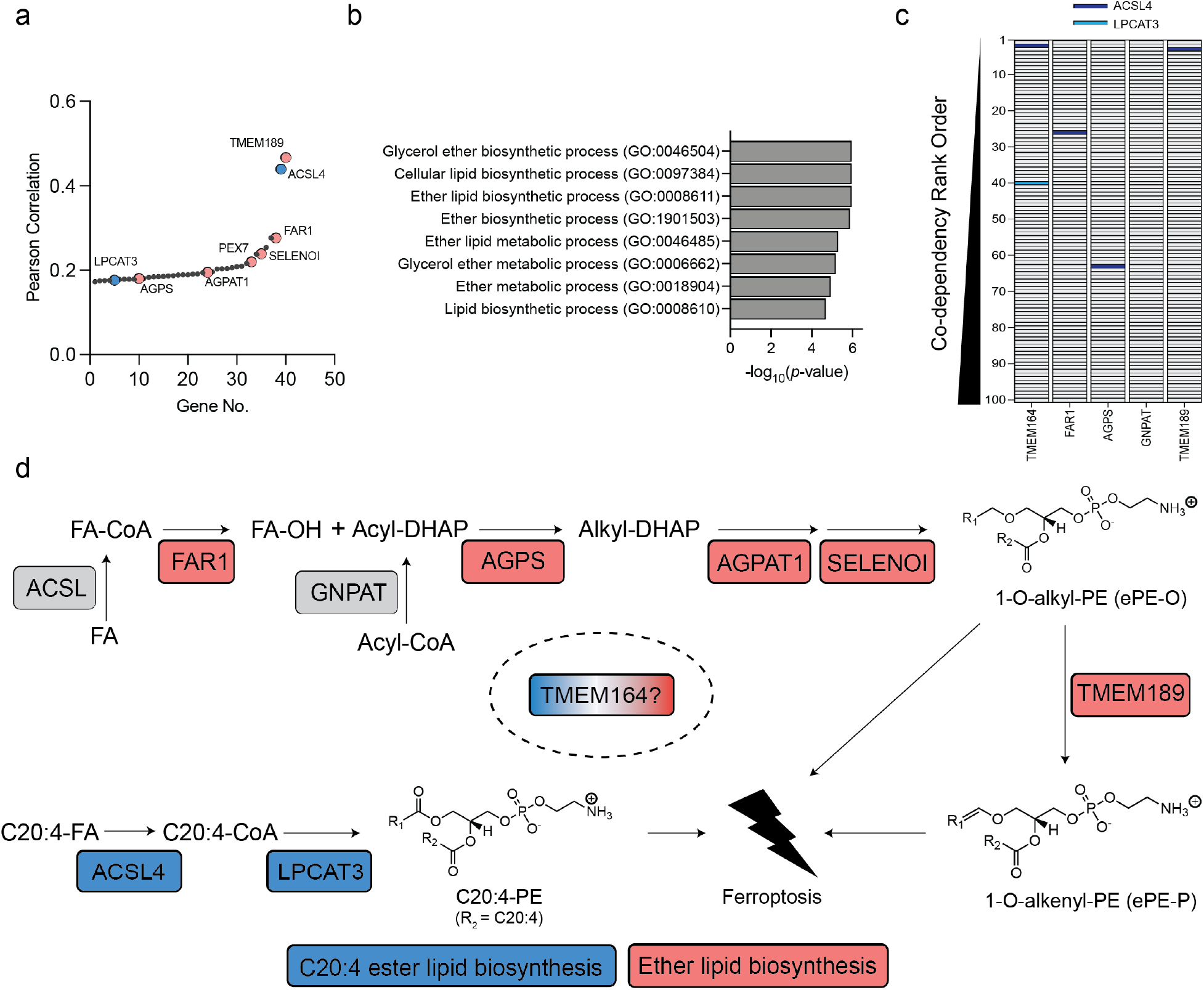
Genome-wide co-dependency screens reveal that TMEM164 has relatedness to PUFA (C20:4) and ether lipid (ePL) metabolic pathways. **a**, Cancer dependency map analysis of TMEM164 showing the genes with the top-40 highest co-dependency scores. Blue and red designate genes involved in C20:4 and ePL metabolism, respectively. **b**, GO enrichment analysis of the top 40 co-dependency genes with TMEm164. **c**, Depiction of the location of C20:4 lipid metabolic genes ACSL4 (red) and LPCAT3 (blue) in the relative rank order of the top 100 co-dependencies for TMEM164 versus established ePL-related genes FAR1, AGPS, GNPAT, and TMEM189. Note that LPCAT3 was only found in the top-100 codependencies for TMEM164. **d**, Metabolic pathway diagrams for C20:4 diester phosphatidylethanolamine (C20:4-PE) and ether PE (ePE) lipids, with C20:4 and ePL enzymes showing co-dependency with TMEM164 highlighted in blue and red, respectively. FA, fatty acid.

### TMEM164 regulates C20:4-ePL production in human cells

We next performed a targeted lipidomic analysis of sgTMEM164 786-O cells using a tandem liquid chromatography mass spectrometry (LC-MS)-based protocol that quantitatively measured ether phosphatidylethanolamine (ePE) lipids, including both acyl chain composition and the presence (plasmenyl, ePE-P) or absence (plasmanyl, ePE-O) of a vinyl ether bond in the sn-1 position (**Extended Data Fig. 2** and **3**). We found that, in comparison to control cells expressing a non-targeted sgRNA (sgCtrl), sgTMEM164 cells displayed major decreases in C20:4 ePEs (**Fig. 2a-c** and **Supplementary Dataset 1**) accompanied by substantial elevations in saturated and monounsaturated ePEs (**Fig. 2a, d, e** and **Supplementary Dataset 1**). These results pointed to a broad acyl-chain remodeling outcome associated with TMEM164 deletion that is reminiscent of the reshuffling of acyl chains in ester PLs that accompanies the genetic or pharmacological disruption of C20:4-preferring ester PL acyltransferases LPCAT3 or MBOAT7^22–25^. One difference in this remodeling profile, however, was the very minor increase in C22:4 ePEs caused by TMEM164 disruption (**Fig. 2d, e** and **Supplementary Dataset 1**), which contrasted with the more dramatic elevation in C22:4 Pes found in LPCAT3-null cells^25,26^. Unlike these ePE changes, the diester PE content of sgTMEM164 cells was largely unaltered (**Fig. 2f**). Reductions in C20:4 ether phosphatidylcholine (ePC) lipids were also observed in sgTMEM164 cells, alongside a more substantial elevation in C20: 4 ester PC lipids (compared to C20:4 ester PE lipids) (**Extended Data Fig. 4** and **Supplementary Dataset 1**).

**Figure 2.**
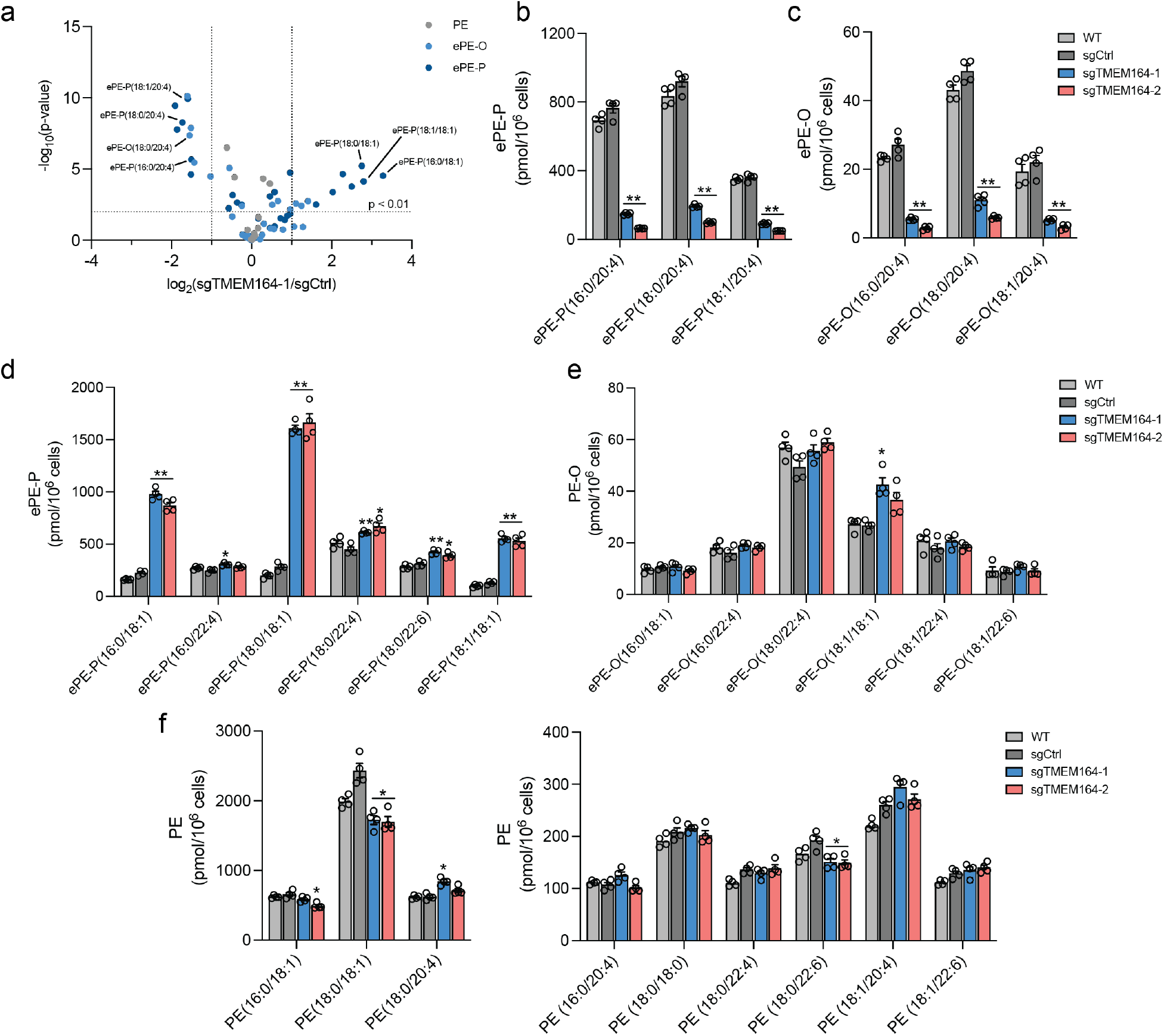
Impairment of C20:4 ePE production in TMEM164-deficient cells. **a**, Volcano plot of targeted lipidomic analysis comparing ether PE (ePE) lipids in sgCtrl and sgTMEM164 786-O-Cas9 cells. Data represent mean values from four independent experiments for sgCtrl and sgTMEM164-1 cells. **b**, **c**, C20:4 ePE-P (**b**) and ePE-O (**c**) lipid measurements in parental (WT), sgCtrl, and sgTMEM164 786-O cells. Two independently generated sgTMEM164 cell populations were analyzed (−1 and −2). **d, e**, Additional ePE-P (**d**) and ePE-O (**e**) lipid measurements in WT, sgCtrl, and sgTMEM164 786-O cells. **f,** Diester PE lipid measurements in WT, sgCtrl, and sgTMEM164 786-O cells. For **b-f**, data represent mean values ± S.E.M from four independent experiments per group. * p < 0.01; ** p < 0.001 (Two-sided Student’s t-test performed relative to sgCtrl cells).

Isotopic tracing experiments provided further evidence that TMEM164 was directly responsible for incorporating C20:4 acyl chains into ePEs. We specifically found that sgTMEM164 cells showed greatly impaired incorporation of deuterated C20:4 [arachidonic acid (AA)-d8] into both ePE-P and ePE-O lipids compared to sgCtrl cells (**Fig. 3a, b** and **Supplementary Dataset 1**). In contrast, the genetic disruption of TMEM164 did not substantially affect the incorporation of AA-d8 into diester PE pools (**Fig. 3c**). Interestingly, we observed much greater incorporation of isotopically labeled monounsaturated fatty acid (C18:1, OA-d9) into ePLs in sgTMEM164 cells (**Fig. 3d** and **Supplementary Dataset 1**), mirroring the increased endogenous concentrations of monounsaturated ePEs observed in these cells (**Fig. 2c, d** and **Supplementary Dataset 1**). This change was also restricted to ePLs, as equivalent amounts of monounsaturated diester PEs were generated in OA-d9-treated sgCtrl and sgTMEM164 cells (**Fig. 3e** and **Supplementary Dataset 1**).

**Figure 3.**
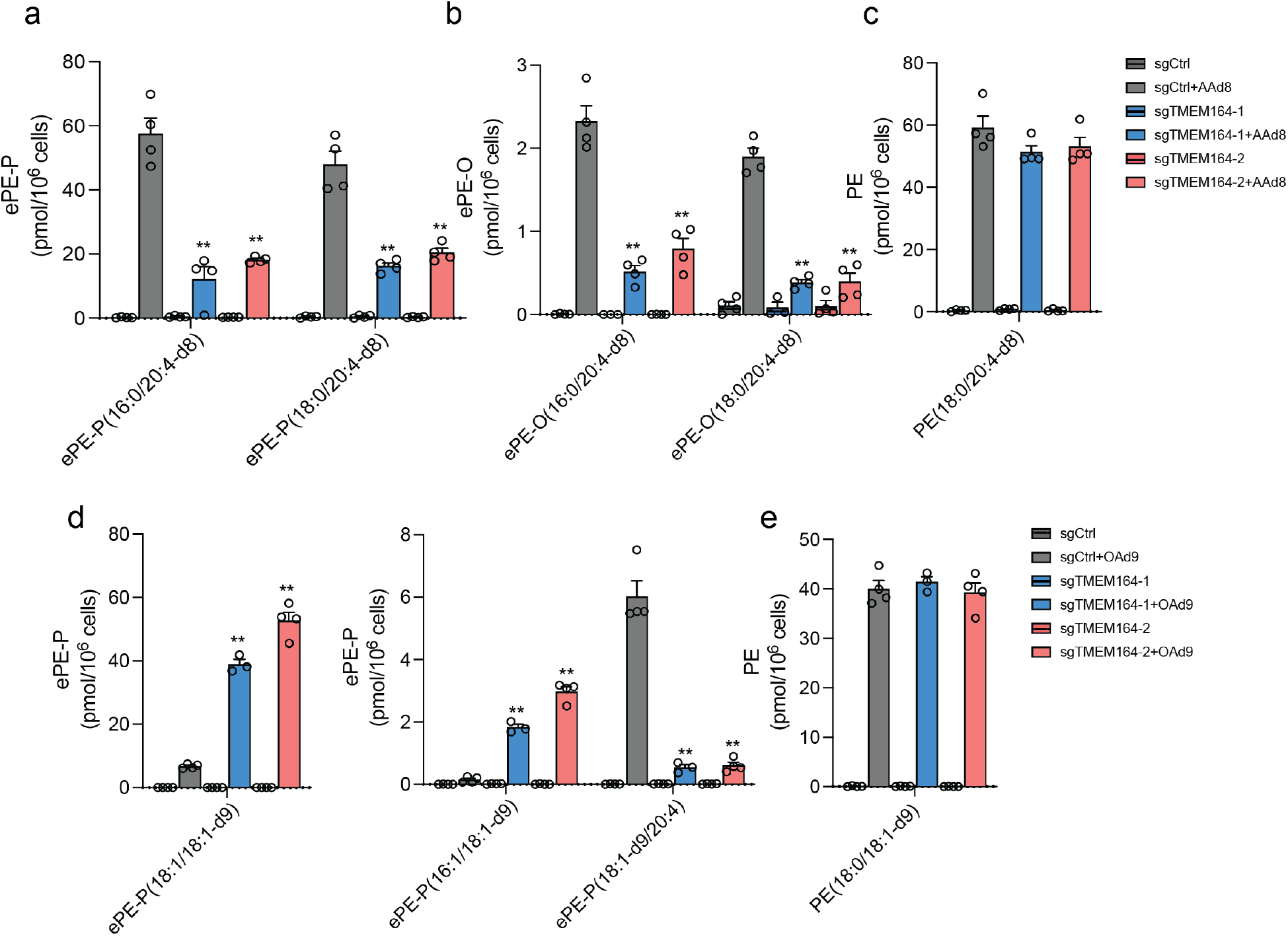
Impaired incorporation of C20:4 (arachidonic acid (AA))-d8 into ePE lipids in TMEM164-deficienct cells. **a-c**, Measurement of AA-d8 incorporation into C20:4 (**a**) ePE-P, (**b**) ePE-O and (**c**) diester PE lipid species in sgCtrl and sgTMEMW4 786-O-Cas9 cells. **d**, **e**, Measurement of oleic acid (OA; C18:1)-d9 incorporation into C18:1 (**d**) ePE-P and (**e**) ester PE lipid species in sgCtrl and sgTMEM164 786-O-Cas9 cells. For **a-e**, cells were treated with 25 μM of AA-d8 or OA-d9 for 4 h prior to lipid measurements. Data represent mean values ± S.E.M from three-four independent experiments per group. ** p < 0.001 (Two-sided Student’s t-test performed relative to sgCtrl cells).

Our lipidomic data, taken together, pointed to a specific function for TMEM164 in regulating C20:4 ePLs that was reminiscent of the catalytic activities of well-established acyltransferases from the MBOAT family (LPCAT3, MBOAT7) involved in the formation of C20:4 diester phospholipids. However, TMEM164 shared no discernible sequence motif or predicted domain homology with the MBOAT acyltransferases. We therefore considered whether TMEM164 might be part of a distinct transmembrane protein/enzyme family.

### Computational evidence that TMEM164 is a transmembrane acyltransferase

In parallel with our experimental characterization of TMEM164 in ePL metabolism, we were engaged in a separate line of inquiry involving integrated deep homology searches and AlphaFold2 modeling^27^ of the AIG1/ADTRP family of animal and fungal multi-pass transmembrane (TM) proteins (PFAM domain family PF04750) that we recently discovered to hydrolyze the FAHFA (fatty acid esters of hydroxylated fatty acids) class of lipids through a likely Thr/His catalytic dyad^28,29^. Iterative PsiBLAST^30^ and HhPRED^31^ searches detected two distant orphan clusters at 10-15% identity: i) YwaF proteins in bacteria and archaea (PF09529), and ii) TMEM164 proteins in animals (PF14808). The derivation of structurally accurate AlphaFold2 models^27^, either from proteome model databases^32^ or built with ColabFold^33^, allowed for a broader structural comparison with DALI^34^ unhindered by low to negligible sequence similarity that, in conjunction with reciprocal homology searches, linked the AIG1/ADTRP, YwaF, and TMEM164 families to additional bacterial/archaeal (YpjA) and protozoa (Hypothetical)) proteins (**Fig. 4a** and **Extended Data Fig. 5a-c**).

**Figure 4.**
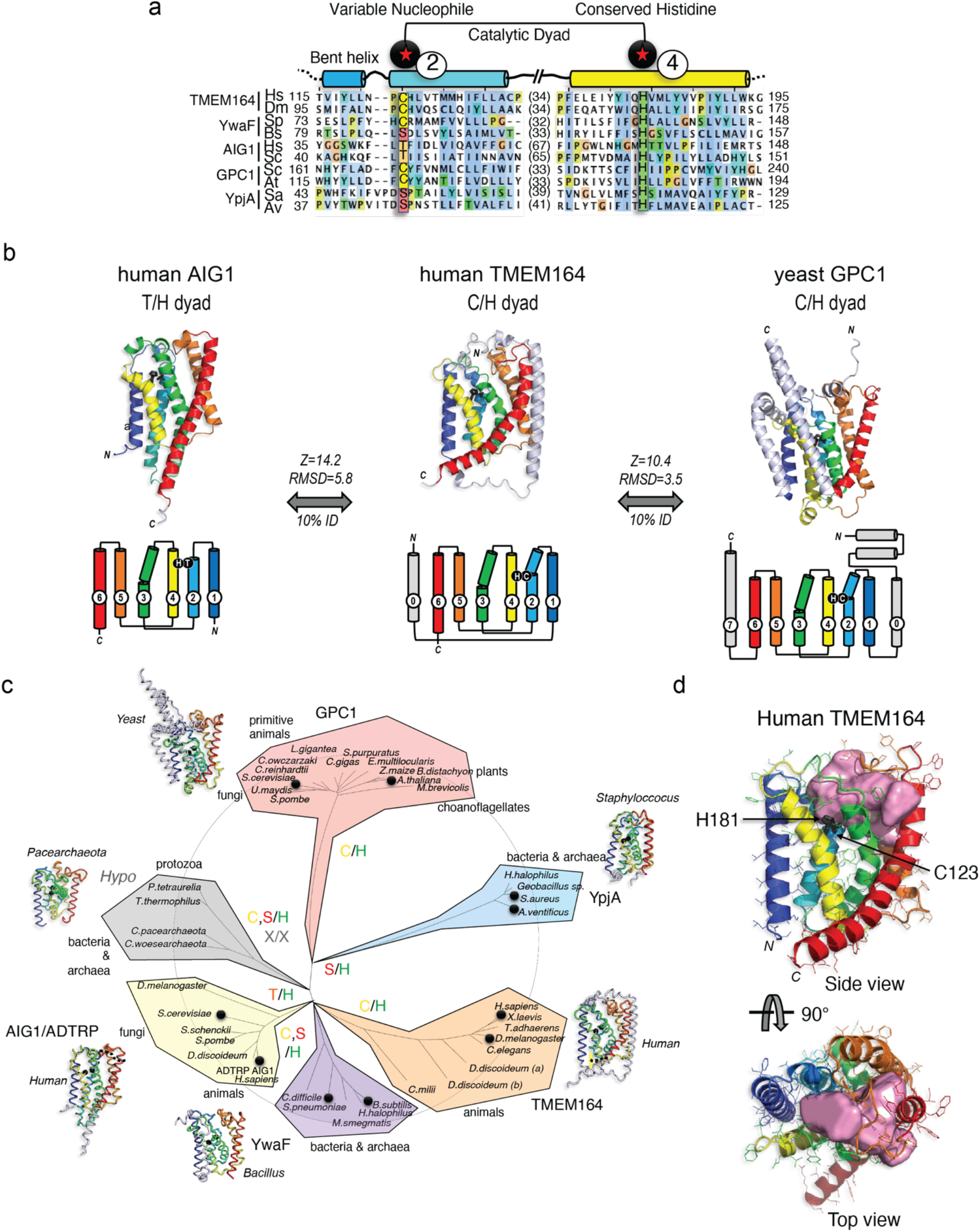
Integrated homology mapping and AlphaFold2 analysis identifies a predicted shared structure between TMEM164 and the AIG1/ADTRP lipid hydrolases. **a**, DALI superposition of the AlphaFold2-predicted structures of human *(H.sapiens;* Hs) and *D. melanogaster* (Dm) TMEM164 reveals a catalytic dyad of C123 and H181 in the human chain (respectively on TM helices 2 and 4 of the conserved 6TM fold) that aligns with the Thr/His dyad of AIG1-class FAHFA hydrolases (from *H.sapiens* and *S. cerevisiae* (Sc)), and defines a superfamily of nucleophile-variable/His-invariant enzymes that share a conserved membrane topology with otherwise sparse sequence identity. Prokaryotic YwaF and YpjA enzymes respectively include *S.pneumoniae* (Sp) and *B.subtilis* (Bs), and *S.aureus* (Sa) and *A.ventificus* (Av) proteins, while GPC1 enzymes are drawn from *S.cerevisiae* and *A.thaliana* (At). See **Extended Data Fig. 5** for a more complete sequence alignment. **b**, Fold and topology comparison of human TMEM164 with human AIG1 and yeast GPC1 enzymes highlighting a 6TM core fold (with helices 1-6 color-ramped to match the topology diagrams; additional helices in grey) with active site Cys or Thr nucleophiles in blue TM2, placed opposite to the invariant His in yellow TM4 (sidechains in black stick format). Z-scores and RMSD values (in Å) for comparison of TMEM164 and AIG1 or GPC1 AlphaFold2 models are provided. **c**, Evolutionary tree of the AIG1/TMEM164/GPC1/YwaF/YpjA (or ATGY) superfamily drawn by DALI-derived fold similarity with variable nucleophilic residues noted in each branch. Black circles mark sequences present in the abridged alignment in **a**. **d**, CavityPlus analysis of the human TMEM164 6TM core showing an internal cavity (with predicted portals to the lipid bilayer) that abuts the C123/H181 catalytic dyad (in black stick form). Figures drawn with Jalview (http://jalview.org) <u>and PyMOL</u> (http://pymol.org).

The AlphaFold2 model showed the preservation of a six transmembrane (6TM) helical core across all of the families (**Fig. 4b, c** and **Extended Data 5d**). Interestingly, however, the equivalent residues to the Thr/His catalytic dyad in the AIG1/ADTRP family were substituted with a conserved Cys/His pair in TMEM164 proteins, while YwaF and YpiA proteins variably displayed Ser, Thr, or Cys nucleophilic residues alongside the invariant His (**Fig. 4a, b** and **Extended Data Fig. 5a, d**). The loose cluster of hypothetical proteins (in bacteria, archaea and protozoa) showed divergent or missing active site residues (**Fig. 4c** and **Extended Data Fig. 5b, c**). The conserved 6TM core of human TMEM164, trained against the UniProt-enlarged AlphaFold2 model databases with the fast structure search engines of RUPEE and FoldSeek^35,36^, collected one additional fold family at less than 10% sequence identity (that had eluded HhPRED queries), centered in fungi, plants, and primitive animals (PF10998), and headlined by the yeast glycerophosphocholine acyltransferase GPC1 (or GPCAT)^37^ (**Fig. 4a-c** and **Extended Data Fig. 5a-d**). Superposition of distant TMEM164 and GPC1 enzymes revealed the shared presence of the predicted Cys/His catalytic dyad in the 6TM core (**Fig. 4a-c**).

The AlphaFold2-predicted structure of human TMEM164 analyzed by CASTp and CavityPlus^38,39^ further revealed an internal cavity with predicted portals to the lipid bilayer and the cytosol abutting the C123/H181 putative catalytic dyad (**Fig. 4d**). CavPHARMER^39^ produced a pharmacophore scaffold within the TMEM164 internal cavity that was compatible with accommodating branched acyl chains of a phospholipid product (**Extended Data Fig. 5e)**.

These structural modeling studies drawing connectivity to other enzymes with experimentally defined roles in lipid metabolism (AIG1 and GPC1 branches), combined with our lipidomic data, pointed to a function for TMEM164 as a cysteine-dependent integral membrane acyltransferase that specifically generates C20:4 ePLs in human cells.

### TMEM164 is an acyltransferase that uses C20:4 PC as an acyl chain donor

In considering candidate substrates for the putative acyltransferase activity of TMEM164, we noted a body literature dating back to the early 1980s describing a transmembrane enzyme activity that transferred the *sn-2* C20:4 acyl chain from PC lipids to lyso-ePL to form C20:4 ePL^15–18^. While this CoA-independent acyltransferase activity (**Fig. 5a**) has been detected in several cell and tissue types^15–18^, it has remained, to date, molecularly uncharacterized. We found that membrane lysates from sgCtrl cells formed C20:4 ePE-P when incubated with deuterated C20:4 PC (and to a lesser extent deuterated C20:4 PE) and lyso-ePE-P, while this activity was ablated in sgTMEM164 cells (**Fig. 5b**). No such PC/PE-dependent acyltransferase activity was observed in sgCtrl membrane lysates incubated with saturated or monounsaturated PC or PE lipids (**Fig. 5c**), consistent with previous studies showing stringent preference for the C20:4 acyl chain^15–18^. Expression of WT-TMEM164, but not a C123A-TMEM164 mutant, restored the C20:4 PC-dependent acyltransferase activity of membrane lysates from sgTMEM164 cells (**Fig. 5d** and **Extended Data Fig. 6a**). We also found that the C20:4 PC-dependent acyltransferase activity of sgCtrl membrane lysates was blocked by the cysteine-directed alkylating agent N-ethylmaleimide (**Extended Data Fig. 6b**), as has been described previously for this activity in other cell types^15^. These results, taken together, indicate that TMEM164 functions as a cysteine-dependent acyltransferase generating C20:4 ePEs from C20:4 PC and lyso-ePE substrates.

**Figure 5.**
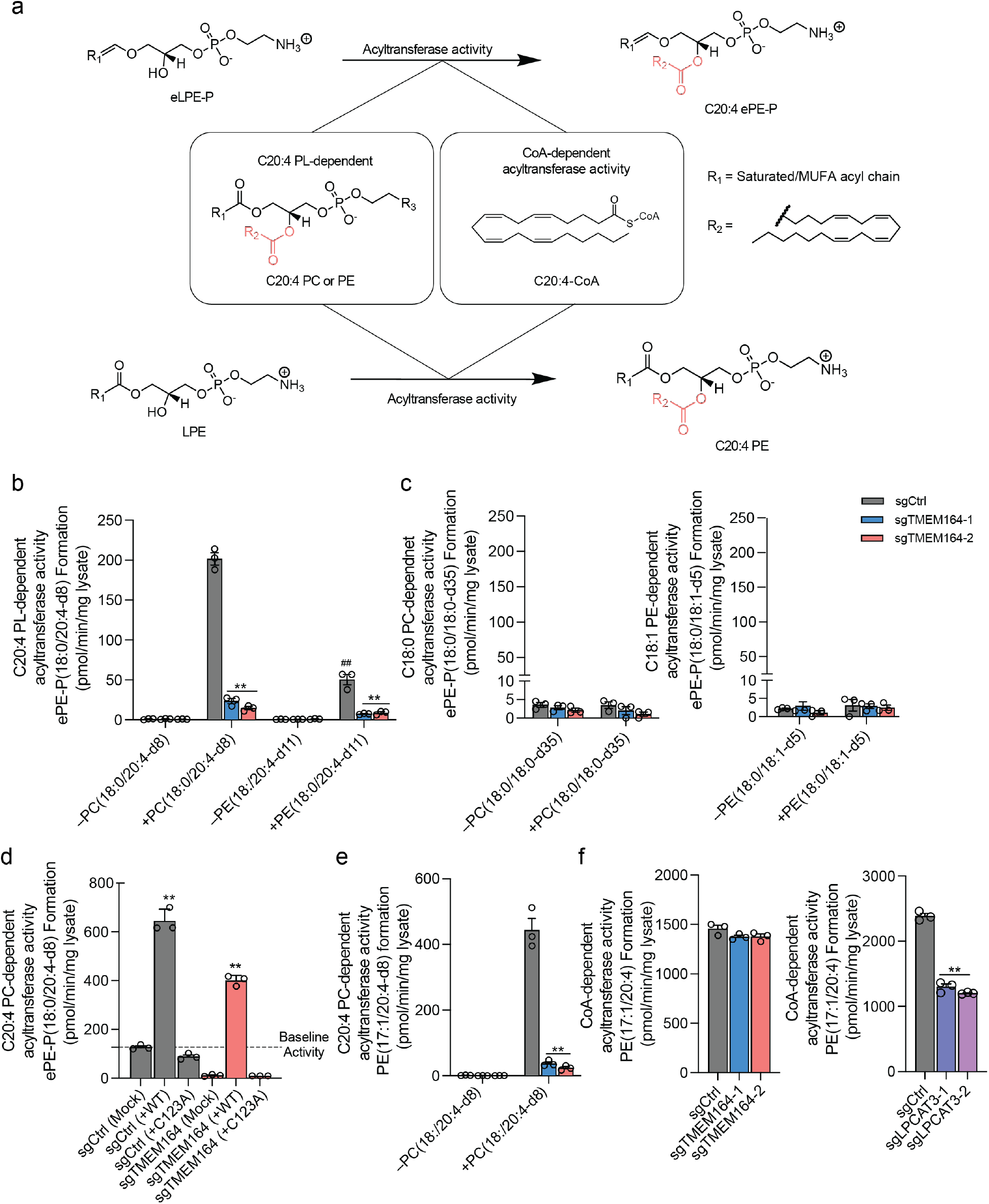
TMEM164 acts as a C20:4-PL-dependent acyltransferase that generates C20:4-ePE lipids. **a**, Diagram of CoA-dependent or C20:4-phospholipid (PL)-dependent acyltransferase reactions responsible for generating C20:4 (e)PE lipids. **b**, Measurement of the formation of ePE-P(18:0/20:4-d8/d11) from diester PC(18:0/20:4-d8) or PE(18:0/20:4-d11) and lyso-ePE-P(18:0) as donor and acceptor substrates, respectively, assayed with membrane lysates from sgCtrl and sgTMEM164 786-O-Cas9 cells. **c**, Measurement of the formation of ePE-P(18:0/18:0-d35/d5) from diester PC(18:0/18:0-d35) or PE(18:0/18:1-d5) and lyso-ePE-P(18:0) as donor and acceptor substrates, respectively, assayed with membrane lysates from sgCtrl and sgTMEM164 786-O-Cas9 cells. **d**, Measurement of the formation of ePE-P(18:0/20:4-d8) from diester PC(18:0/20:4-d8) and lyso-ePE-P(18:0) from membrane lysates of sgCtrl and sgTMEM164-2 786-O-Cas9 cells engineered to recombinantly express FLAG-tagged WT-TMEM164 or a C123A-TMEM164 mutant. The cell lysates containing WT-TMEM164 were diluted 1:1 with mock lysate to normalize expression to the C123A-TMEM164 mutant (see **Extended Data Fig. 6a**). **e**, Measurement of the formation of diester PE(17:1/20:4-d8) from diester PC(18:0/20:4-d8) and LPE(17:1) as donor and acceptor substrates, respectively, assayed with membrane lysates from sgCtrl and sgTMEM164 786-O-Cas9 cells. **f**, Measurement of the formation of diester PE(17:1/20:4) using C20:4-CoA and LPE(17:1) as donor and acceptor substrates, respectively, assayed with membrane lysates from sgCtrl, sgTMEM164, and sgLPCAT3 786-O cells. For **b-f**, membrane lysates (1 μg/μL) were treated with 50 μM each of donor and acceptor substrates for 1 h at 37° C prior to analysis. Data represent mean values ± S.E.M from three independent experiments per group. ** p < 0.001 (Two-sided Student’s t-test performed relative to sgCtrl cells). For **b**, ## p < 0.001 [Two-sided Student’s t-test performed relative to sgCtrl: +PC(18:0/20:4-d8)].

While our findings supported that TMEM164 represents the historically described C20:4 PC-dependent acyltransferase activity responsible for C20:4 ePL formation, we wondered whether this protein might also utilize C20:4 CoA as a substrate. This question initially proved challenging to address, as neither isotopicially labeled C20:4 CoA nor nonnatural acyl chain (e.g., C17 acyl chain) variants of lyso-ePLs are commercially available. We observed, however, that sgCtrl, but not sgTMEM164 membrane lysates, also supported the formation of C20:4 diester PE using deuterated C20:4 PC and C17:1 lyso-PE as acyl chain donor and acceptor lipids, respectively (**Fig. 5e**). This discovery allowed for subsequent testing of CoA-dependent acyltransferase activity for TMEM164 by incubating sgCtrl and sgTMEM164 membrane lysates with C20:4 CoA and C17:1 lyso-PE and measuring production of the unnatural lipid C20:4/C17:1 PE. Using this assay, we found that sgCtrl and sgTMEM164 lysates showed equivalent C20:4 CoA-dependent acyltransferase activity, which was instead substantially reduced in membrane lysates from sgLPCAT3 cells (**Fig. 5f**). On the other hand, sgLPCAT3 membrane lysates exhibited unaltered C20:4 PC-dependent acyltransferase activity using both C17:1 lyso-PE and C18:0 lyso-ePE-P as acceptor substrates (**Extended Data Fig. 6c, d)**, indicating that LPCAT3 and TMEM164 serve as exclusively C20:4 CoA- and C20:4 PC-dependent acyltransferases, respectively.

### TMEM164-C123Y cells show reduced C20:4 ePLs and ferroptotic sensitivity

We recombinantly expressed WT-TMEM164 and several C123X point mutants in HEK293T cells and found that all of the C123 mutants exhibited negligible acyltransferase activity (**Fig. 6a, b**). Several of the C123X mutants expressed at ~50% lower levels than WT-TMEM164 (e.g., C123A, C123S, and C123V), and the C123R mutant was not detectable (**Fig. 6a**). The C123Y mutant, however, expressed at similar levels to WT-TMEM164 (**Fig. 6a**), which was a noteworthy finding because base editing methods using a cytidine deaminase convert cysteine residues to tyrosine^40^. We therefore used cytosine base editing to create a TMEM164-C123Y knock-in population of 786-O cells (**Extended Data Fig. 7a**) and compared their properties to 786-O cells treated with a sgRNA targeting a non-human (GFP) control gene (sgCtrl). We found that the TMEM164-C123Y cells displayed substantial reductions in C20:4 PC-dependent acyltransferase activity (**Fig. 6c**) and C20:4 ePE lipids (**Fig. 6d**, **Extended Data Fig. 7b** and **Supplementary Dataset 1**) with concomitant elevations in saturated/monounsaturated ePE lipids (**Fig. 6e**) and unchanged diester PE lipids (**Extended Data Fig. 7c** and **Supplementary Dataset 1**). Finally, the TMEM164-C123Y knock-in 786-O cells also displayed reduced sensitivity to GPX4 inhibitor-induced ferroptosis (**Fig. 6f** and **Extended Data Fig. 7d**). Considering that the biochemical and cellular changes observed in the TMEM164-C123Y knock-in cells generally matched those found in TMEM164-null (sgTMEM164) cells (see **Extended Data Fig. 1c, d** and **Fig. 2**), these results indicate that the contributions of TMEM164 to ePL metabolism and ferroptosis depend on the C123-mediated acyltransferase activity of this protein.

**Figure 6.**
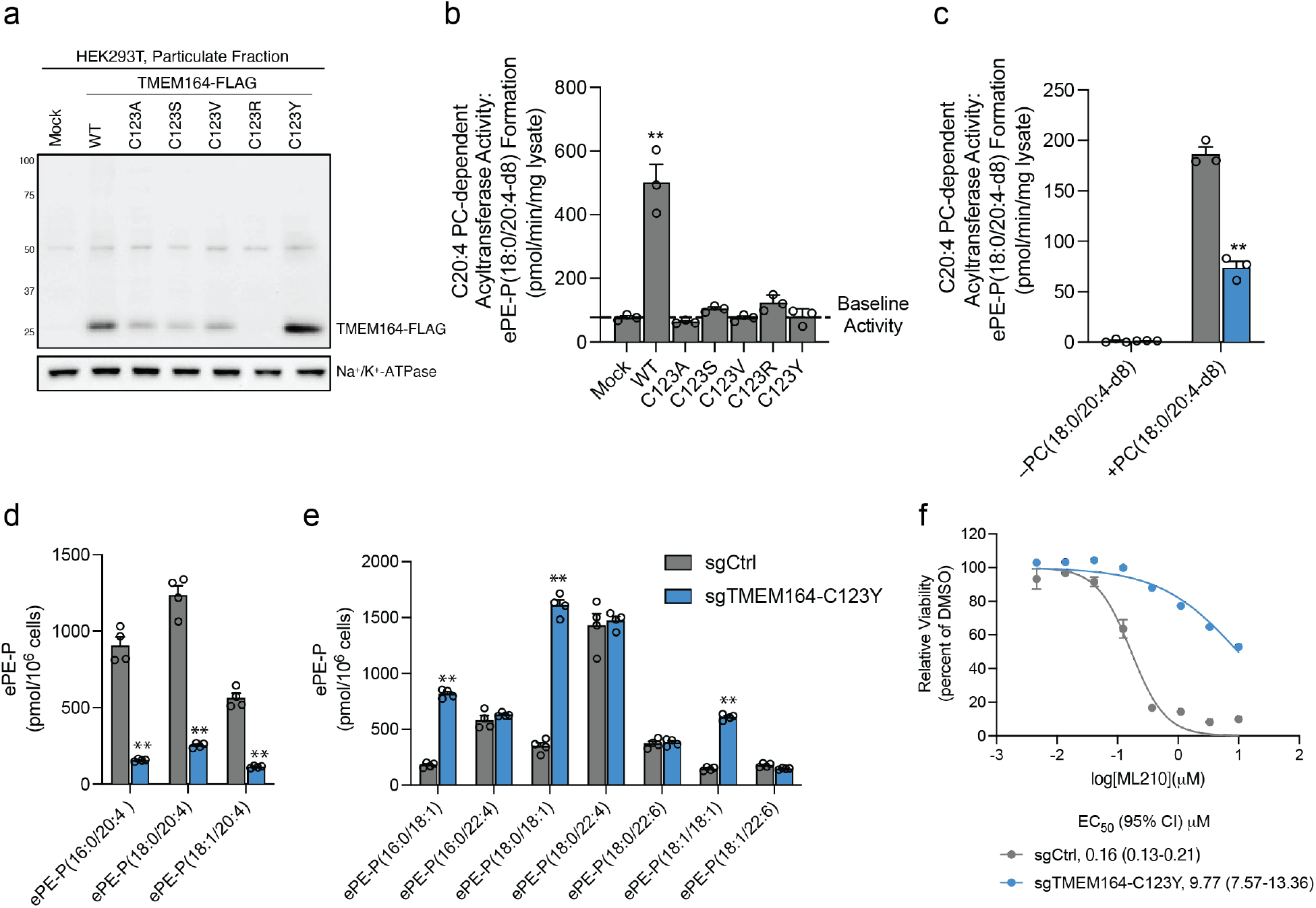
Regulation of ePL content and ferroptotic sensitivity by TMEM164 requires C123. **a,** Western blot of membrane lysates from HEK293T cells expressing C-terminally FLAG-tagged WT-TMEM164 or C123X-TMEM164 mutants. **b, c,** Measurement of the formation of ePE-P(18:0/20:4-d8) from diester PC(18:0/20:4-d8) (50 μM) and lyso-e-PE-P(18:0) (50 μM) as substrates assayed with membrane lysates (1 μg/uL, 1 h incubation at 37° C) from (**b**) HEK293T cells expressing the indicated TMEM164 proteins or (**c**) sgCtrl and sgTMEM164-C123Y based-edited 786-O cell populations. **d, e**, Measurement of C20:4 ePE-P (**d**) and additional ePE-P (**e**) lipids in sgCtrl and sgTMEM164-C123Y base-edited 786-O cell populations. **f,** Viability of sgCtrl and sgTMEM164-C123Y base-edited 786-O cell populations treated with the indicated concentrations of ML210 measured at 24 h post-treatment. For **b-e**, data represent mean values ± S.E.M from three-four independent experiments per group. *p < 0.01, ** p < 0.001 (Two-sided Student’s t-test performed relative to sgCtrl cells). For **f**, Data represent mean values ± S.E.M. from two independent experiments with two-three technical replicates per experiment.

## Discussion

The molecular characterization of orphan enzyme activities has benefited from several recent advances, including – i) complete genome sequences of diverse organisms, which enables deep homology mapping of large protein families^41,42^; ii) protein structure prediction programs, such as AlphaFold, which facilitate the discovery of shared folds for very distally related proteins^27^; and iii) large-scale gene essentiality screens across many hundreds of cell lines, which can assign genes to a common biochemical pathway based on correlated dependency profiles^9,19^. Here, we have made use of each of these approaches to determine that TMEM164, a previously uncharacterized integral membrane protein implicated in the regulation of ferroptosis^8,20,21^, acts as an acyltransferase that transfers C20:4 acyl chains from PC to lyso-ePLs to generate C20:4-ePLs – a biochemical reaction first described nearly thirty years ago in the literature^15–18^, but for which the responsible enzyme has remained unknown. We provide evidence that a conserved cysteine (C123) is required for the acyltransferase activity of TMEM164. This cysteine, along with a conserved histidine (H181) also align based on sequence and structural homology predictions with the putative Thr/His catalytic dyad of the AIG1/ADTRP family of lipid hydrolases. Based on this information, we posit that C123 serves as a catalytic nucleophile of TMEM164, first reacting with C20:4 PC to form an acyl-enzyme intermediate, followed by transfer of this C20:4 acyl group to lyso-ePLs to form ePLs (**Extended Data Fig. 8**). Such an acyltransferase mechanism involving acyl-donating PC substrates and presumed protein-bound acyl-enzyme intermediates has precedent in other lipid metabolic pathways, including those that form cholesterol esters^43^ and N-acyl phosphatidylethanolamines^44^. Studies to date with the distantly related enzyme GPC1 indicate that it uses acyl-CoA^37^, suggesting that members of the AIG1/TMEM164/GPC1/YwaF/YpjA (or ATGY) enzyme family may use diverse acyl-donating substrates.

The geometry of the minimal catalytic dyad predicted for the TMEM164 and AIG1/ADTRP enzymes resembles part of the Cys/His/Asn and Ser/His/Asp catalytic triads in papain cysteine proteases and trypsin proteases, respectively^45^. The trypsin superfamily further encompasses viral enzymes evolved to use Cys/His/Asp triads, and likewise, human trypsins can function with Cys or Thr nucleophiles^46–48^, attesting to the plasticity of nucleophile catalysis in a conserved fold setting. These hydrolytic triads can be further whittled to a nucleophile/His dyad—accompanied by an ‘oxyanion hole’ of H-bond donors—grafted into non-protease scaffolds, suggesting a minimal origin for functional esterase active sites^49^.The nucleophilic promiscuity of the ATGY enzyme family appears to represent a particularly provocative example of evolutionary (re)invention of a catalytic engine married to an integral membrane fold suited to different steps in lipid metabolism^50^.

The remarkable specificity displayed by TMEM164 for transferring C20:4 acyl chains suggests that this enzyme has a specialized role in regulating PUFA ePL content, which is reminiscent of the roles that enzymes like LPCAT3 and MBOAT7 play in C20:4 phospholipid metabolism^22–25^. In each case, when the C20:4 acyltransferase is genetically disrupted, the decreases in direct C20:4 (e)PL products is accompanied by increases in (monoun)saturated (e)PLs, which may reflect an acyl-chain remodeling outcome that may serve to maintain a consistent amount of total e(PL) in the cell. One peculiar feature of the TMEM164-disrupted lipid profile was the substantial elevations in C20:4 PCs, which we interpret to indicate that TMEM164 not only regulates basal concentrations of its enzymatic products (C20:4 ePLs), but also the acyl-chain donor substrate C20:4 PC. Despite this elevation in C20:4 PC, the net functional outcome of the lipid changes caused by TMEM164 disruption was highly protective in ferroptotic assays. These data support recent studies indicating a particularly important role for PUFA ether lipids and ePEs, in particular, over diester PC lipids in regulating ferroptosis sensitivity of human cells^7,51^.

Projecting forward, the discovery that TMEM164 is a specialized acyltransferase responsible for regulating C20:4 ePL content in human cells raises several important questions. For instance, we found that TMEM164 also accepted ester lyso-PLs as substrates, but diester PLs were generally unperturbed in TMEM164-disrupted cells. The diester C20:4 PL content was instead regulated by LPCAT3. Does this reflect distinct subcellular localizations for LPCAT3 and TMEM164 that restrict their respective activities to specific lipid substrate classes in the cell? Our efforts to date to analyze TMEM164 with commercially available antibodies have been largely unsuccessful, as we have only been able to detect a weak signal for recombinantly expressed TMEM164 in HEK293T cell membrane lysate and have not yet been able to visualize endogenous TMEM164 in cell lines like 786-O (**Extended Data Fig. 9**). Future efforts to generate higher quality antibodies capable of detecting endogenous TMEM164 are therefore warranted. The presence of a conserved cysteine required for TMEM164 activity also suggests the potential to create covalent inhibitors of the enzyme, which may have value as chemical probes and drugs to suppress ferroptosis. Searches of our legacy chemical proteomic datasets have revealed sparse coverage of TMEM164^52,53^ including the tryptic peptide containing C123, which may reflect low abundance of this integral membrane protein or limited reactivity with the chemical probes evaluated so far. When contemplating potential functions for TMEM164 and C20:4 ePLs beyond ferroptosis, we call attention to the provocative Dependency Map profiles shared by TMEM164, TMEM189, and ACSL4, where subsets of hematological cancer cell lines show impaired growth when the genes encoding these proteins are deleted (https://depmap.org/). Previous studies have also demonstrated that certain cancer cells express heightened ePL content^54–57^. Future lipidomic analyses of TMEM164-dependent cancer cells may accordingly be illuminating. In summary, our work, combined with recent functional assignments of TMEM189 as a plasmanylethanolamine desaturase^9–11^, has furnished a near-complete metabolic pathway for producing C20:4-ePLs, a lipid class with key contributions to cell biological processes such as ferroptosis.

## Acknowledgements

This work was supported by the NIH (CA231991) and Lundbeck. We are grateful to J. Blankman, H. Reardon, and M. Niphakis for helpful discussions throughout the performance of this research.

## Author contributions

A.R., T.W., J.F.B, and B.F.C. conceived of the project and wrote the manuscript. A.R. and T.W. generated genetically engineered cell lines and performed lipidomics, biochemical assays and western blotting experiments. J.F.B. performed computational structural analysis studies linking TMEM164 to AIG1/ADTRP family of enzymes. H.L. assisted with targeted genomic sequencing and data analysis.

## Data Availability.

Data supporting the findings of this work are available within the paper and its Supplementary Information. The datasets, constructs, and cell models generated and analyzed during the current study are available from the corresponding author upon request.

---

**Extended Data Figure 1.**
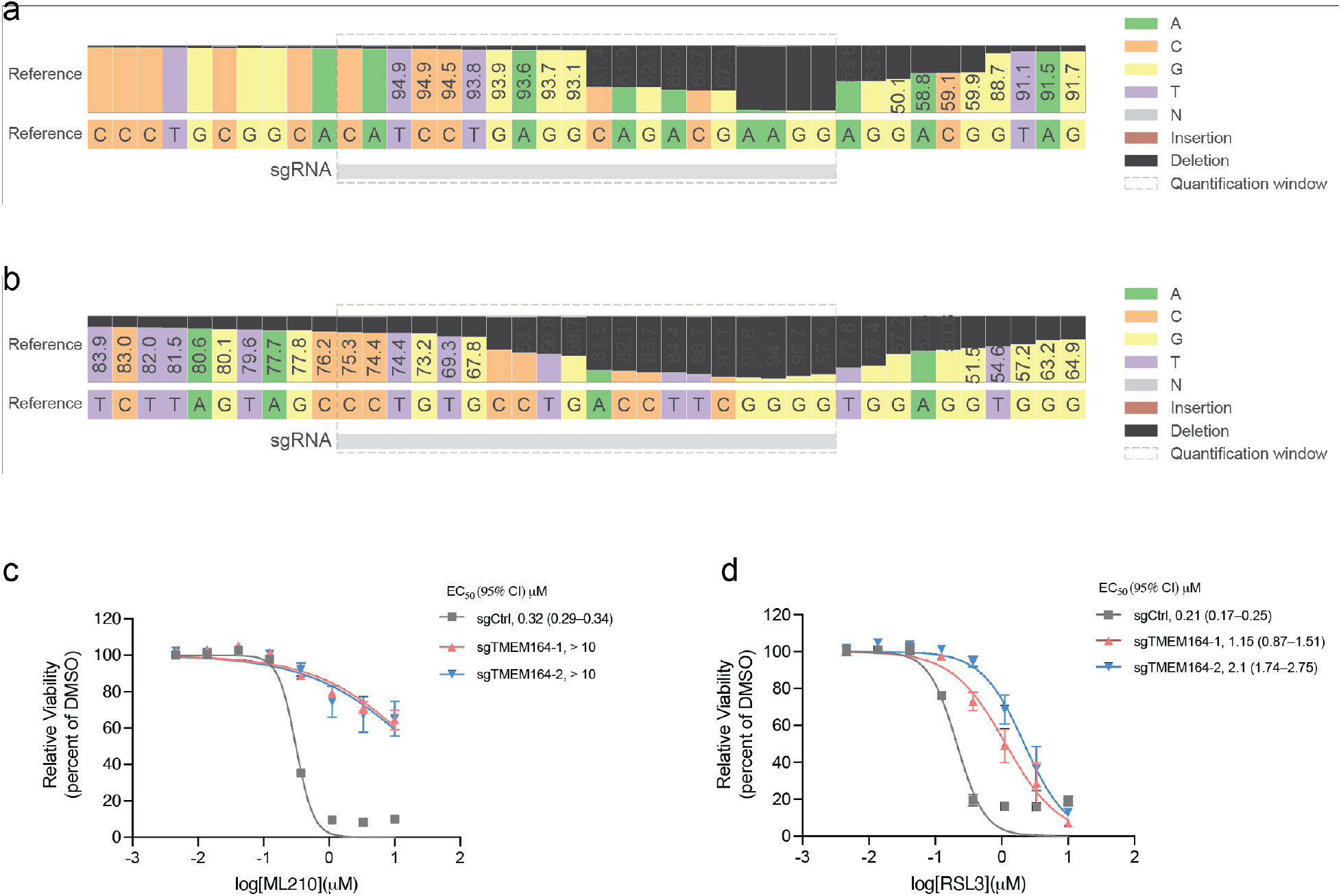
Characterization of sgTMEM164 786-O-Cas9 cells. **a, b,** Confirmation of genetic alterations in 786-O-Cas9 cells at sgRNA sites targeted with (**a**) sgTMEM164-1 and (**b**) sgTMEM164-2 by next-generation sequencing analysis. **c, d,** Genetic deletion of TMEM164 in 786-O-Cas9 cells results in reduced sensitivity to ferroptosis induced by the GPX4 inhibitors ML210 (**c**) and RSL3 (**d**) measured at 24 h post-treatment with GPX4 inhibitors. Data represent mean values ± S.E.M. from two independent experiments with three technical replicates per experiment.

**Extended Data Figure 2.**
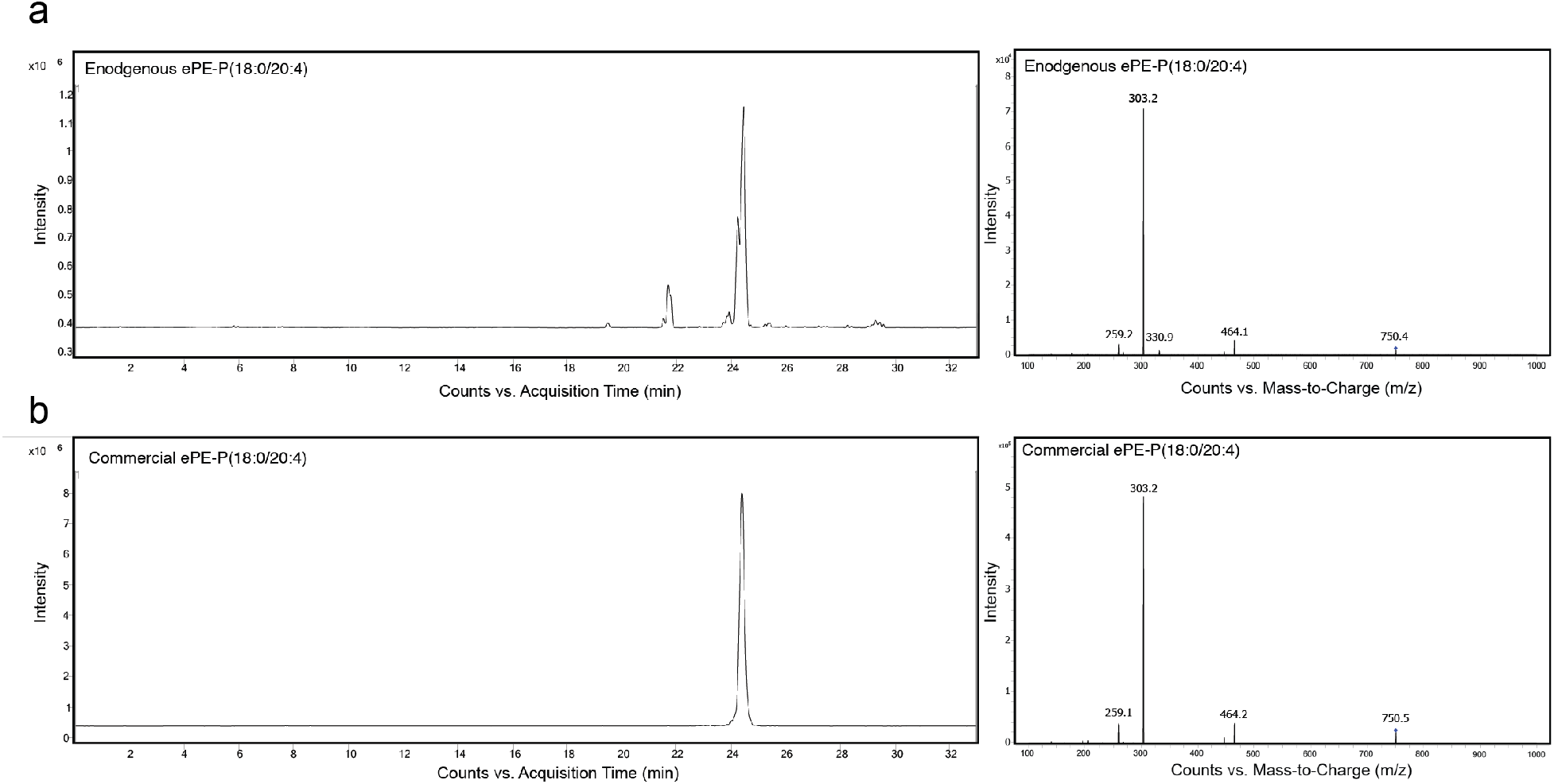
Targeted lipidomic (LC-MS/MS) methods for measuring ePLs. **a, b,** Detection of endogenous ePE-P(18:0/20:4) in 786-O-Cas9 cells by LC-MS/MS (**a**) and comparison to commercial ePE-P(18:0/20:4) standard (**b**). Left traces show extracted ion chromatogram of a feature corresponding to ePE-P(18:0/20:4) (750.5 → 303.3). Right traces show MS2 spectra for ePE-P(18:0/20:4).

**Extended Data Figure 3.**
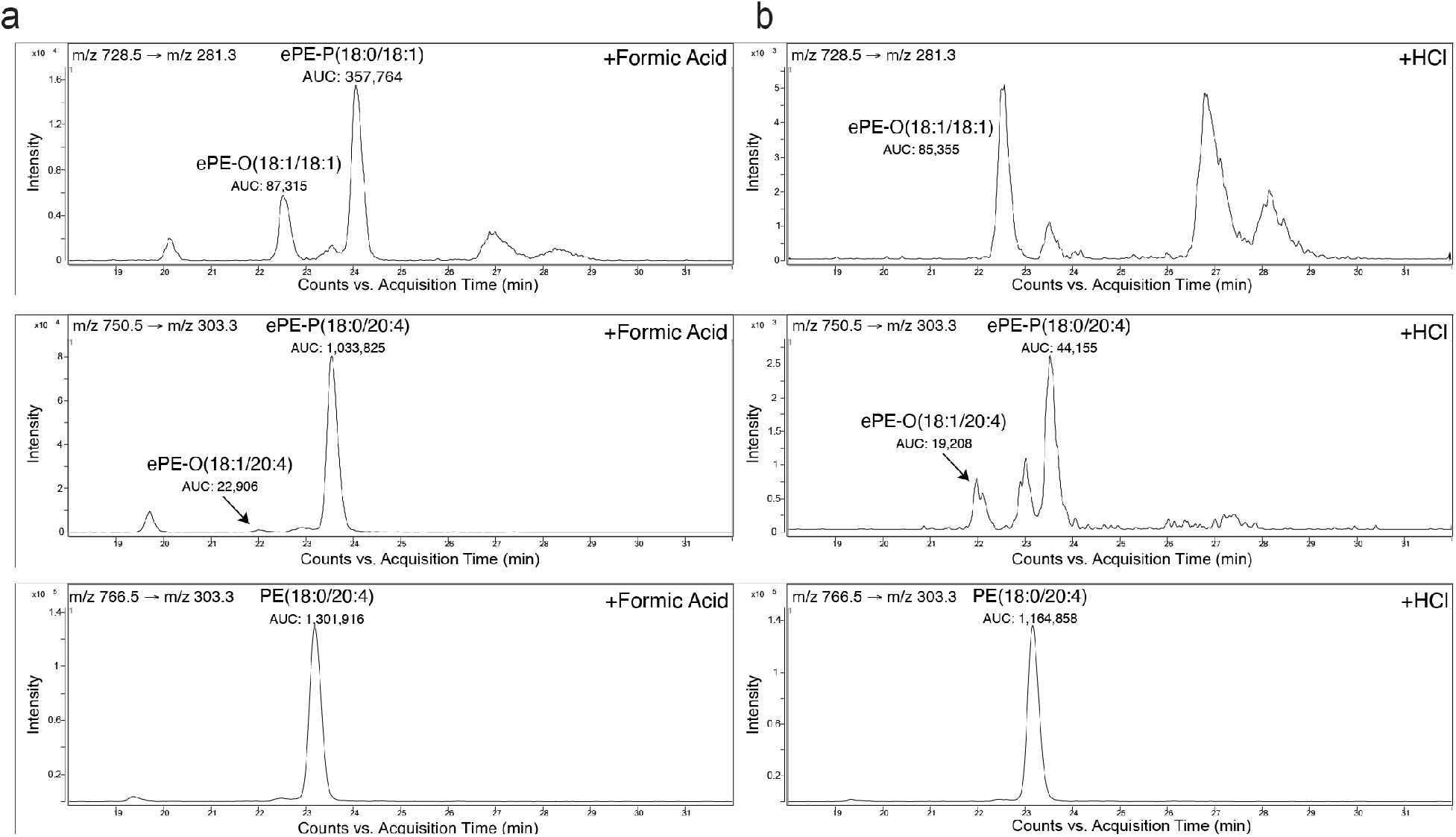
Targeted lipidomic (LC-MS/MS) methods for measuring ePLs. **a,b,** Targeted LC-MS/MS analysis of isobaric ePE-O and ePE-P species performed generally as described^11,58^ revealing that they are separated by approximately 1.5 min difference in retention time. Treatment of lipid extracts from 786-O-Cas9 cells with 10% (v/v) formic acid (**a**) or HCl (3N) (**b**) for 30 min at 37°C reveals ePE-P elutes after ePE-O as evidenced by a significant reduction in the ePE-P peak following HCl treatment. AUC, area under the curve.

**Extended Data Figure 4.**
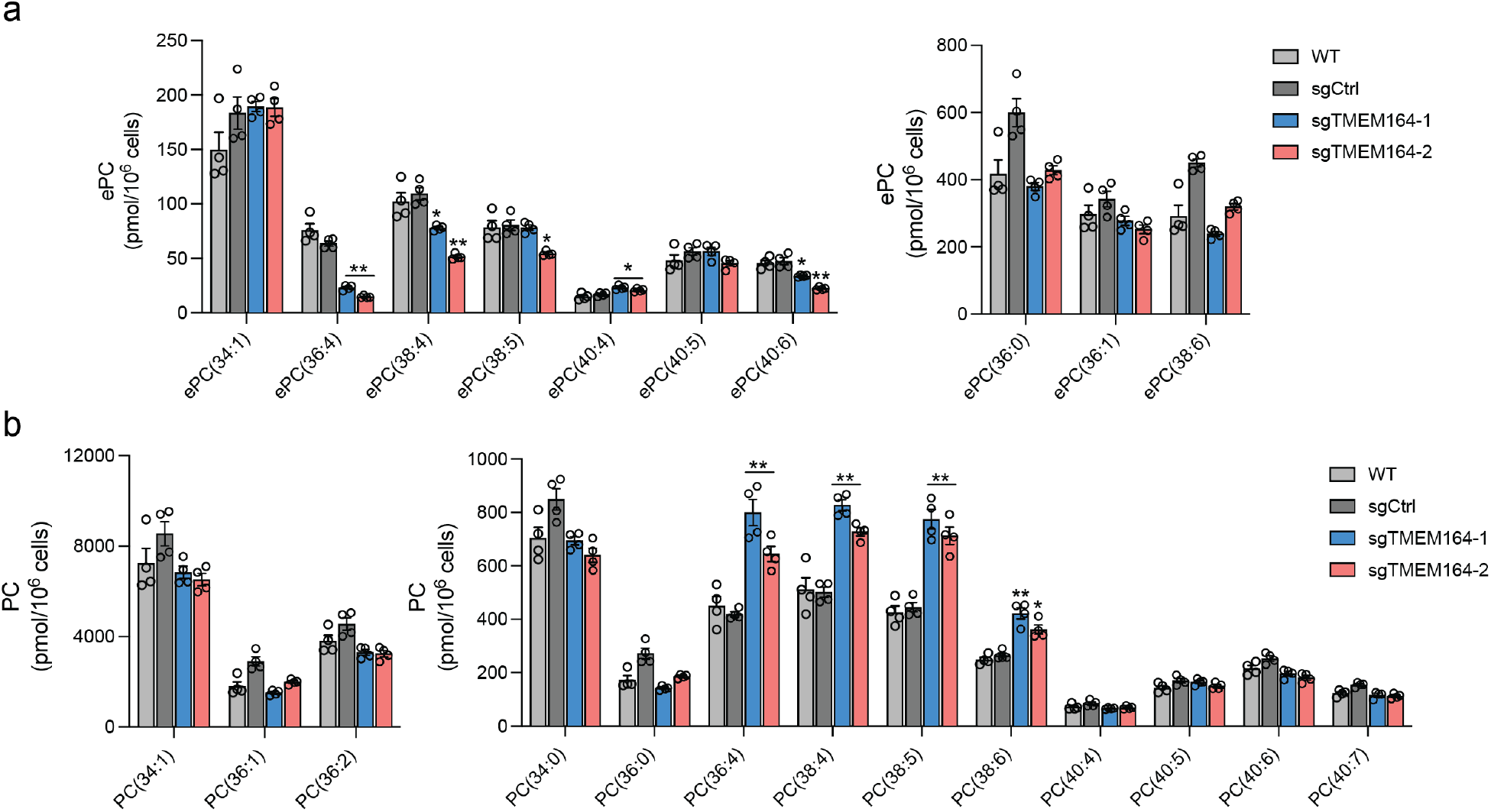
Alterations in ePC and diester PC production in TMEM164-deficient cells. **a, b** ePC (**a**) and PC (**b**) lipid measurements in sgCtrl and sgTMEM164 786-O-Cas9 cells. For **a**, as described in previous protocols^7^, we did not distinguish between ePC-O and ePC-P lipids. Data represent mean values ± S.E.M from four independent experiments per group. * p <0.01; ** p < 0.001 (Two-sided Student’s t-test performed relative to sgCtrl cells; only shown for lipids where sgCtrl and parental (WT) data did not differ).

**Extended Data Figure 5.**
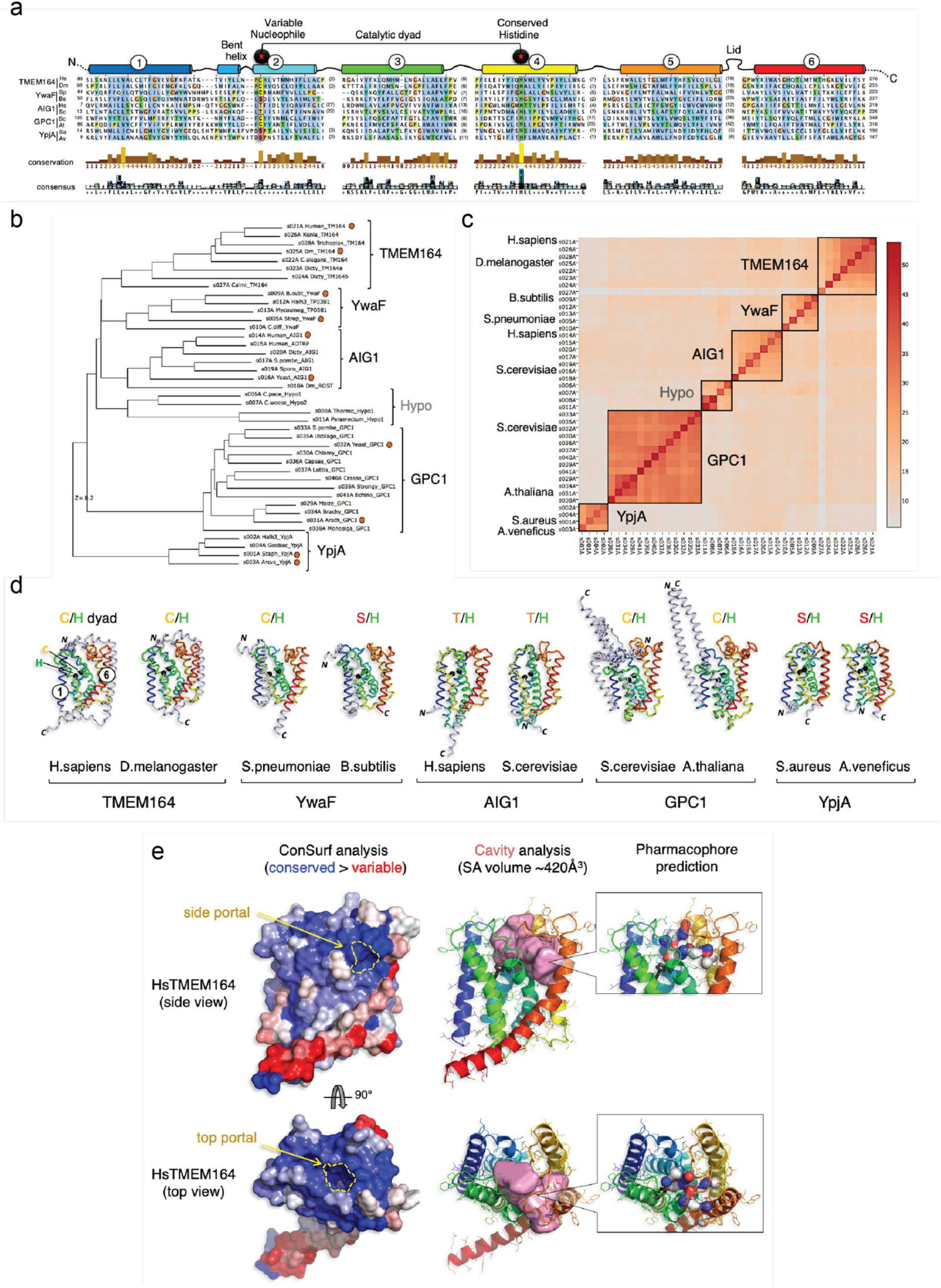
Integrated homology mapping and AlphaFold2 analysis identifies a predicted shared structure between TMEM164 and the AIG1/ADTRP lipid hydrolases. **a**, Structure-based sequence alignment by DALI of AlphaFold2 models retrieved from the AlphaFold database (http://alphafold.ebi.ac.uk) of the H. sapiens (UniProt sequence Q5U3C3) and D. melanogaster (UniProt Q7JRB2) TMEM164 proteins, with representative proteins from the AIG1/ADTRP family (H. sapiens (Q9NVV5) and S.cerevisiae (P38842), from prokaryotic YwaF (S.pneumoniae (Q8CYG1) and B.subtilis (P25149) and YpjA proteins (S.aureus (Q2FYH4) and A.venificus (F2KNL0), and GPC1 enzymes from S.cerevisiae (P48236) and A.thaliana (Q9FJB4). In some cases, models were built anew with the ColabFold implementation of AlphaFold2 (http://github.com/sokrypton/ColabFold). The alignment was displayed and analyzed with Jalview (http://jalview.org), showing the six blocks of conserved structure that correspond to TM helices 1-6 of the TMEM164 fold, color-ramped from blue (TM1) to red (TM6). Length and sequence-variable gaps in the alignment that correspond to loops in the superposed folds were abbreviated by residue stretches in parentheses. The positions of the variable nucleophile and invariant His residues (Cys123 and His181 in human TMEM164, respectively) are boxed and labeled. **b**, DALI-derived superpositions provide a measure of structural relatedness for the six main branches of the enzyme superfamily displayed in the phylogenetic hypertree in **Fig. 4c** (http://kinase.com/tools/HyperTree.html), with the selected chains aligned in panel **a** noted by red circles. Z-score of 8.2 is provided for the most distant branch (YpjA). **c**, The matrix of DALI Z-scores displays the clusters of more closely related structures from all-against-all superpositions of the AlphaFold2-derived structures performed with the DALI server (http://ekhidna2.biocenter.helsinki.fi), leading to reconstitution of the six branches of the enzyme superfamily. Five of these six fold clusters have corresponding sequence alignments that can be retrieved from the PFAM database (http://pfam.xfam.org) for TMEM164 (PF14808), AIG1/ADTRP (PF04750), GPC1 (PF10998), YwaF (PF09529) and YpjA (PF7187). **d**, Tube representations of the aligned AlphaFold2 models showing the positions of the catalytic dyad with black Ca spheres and the preservation of the core 6TM fold (color-ramped from blue TM1 to red TM6) with N-or C-terminal extensions drawn in grey. Figures composed with PyMOL (http://pymol.org). **e**, Human TMEM164 structural model was processed by the ConSurf server (http://consurf.tau.ac.il) to map the sequence conservation profile of its underlying sequence family to its 3D fold surface color-ramped from blue (conserved) to red (variable) patches. For side and top views, the ConSurf surfaces are aligned to the ribbon model of TMEM164 embedded with the CavityPlus-calculated volume in pink (server at http://pkumdl.cn), showing the positions of predicted portals to the lipid bilayer. In addition, the CavPharmer tool at CavityPlus was used to predict a pharmacophore skeleton for a prospective ligand (which could be substrates or product of the enzymatic reaction) for TMEM164, showing a shape and chemical nature in agreement with its predicted function as a Arachidonate-preferring Lyso-Plasmalogen Acyltransferase.

**Extended Data Figure 6.**
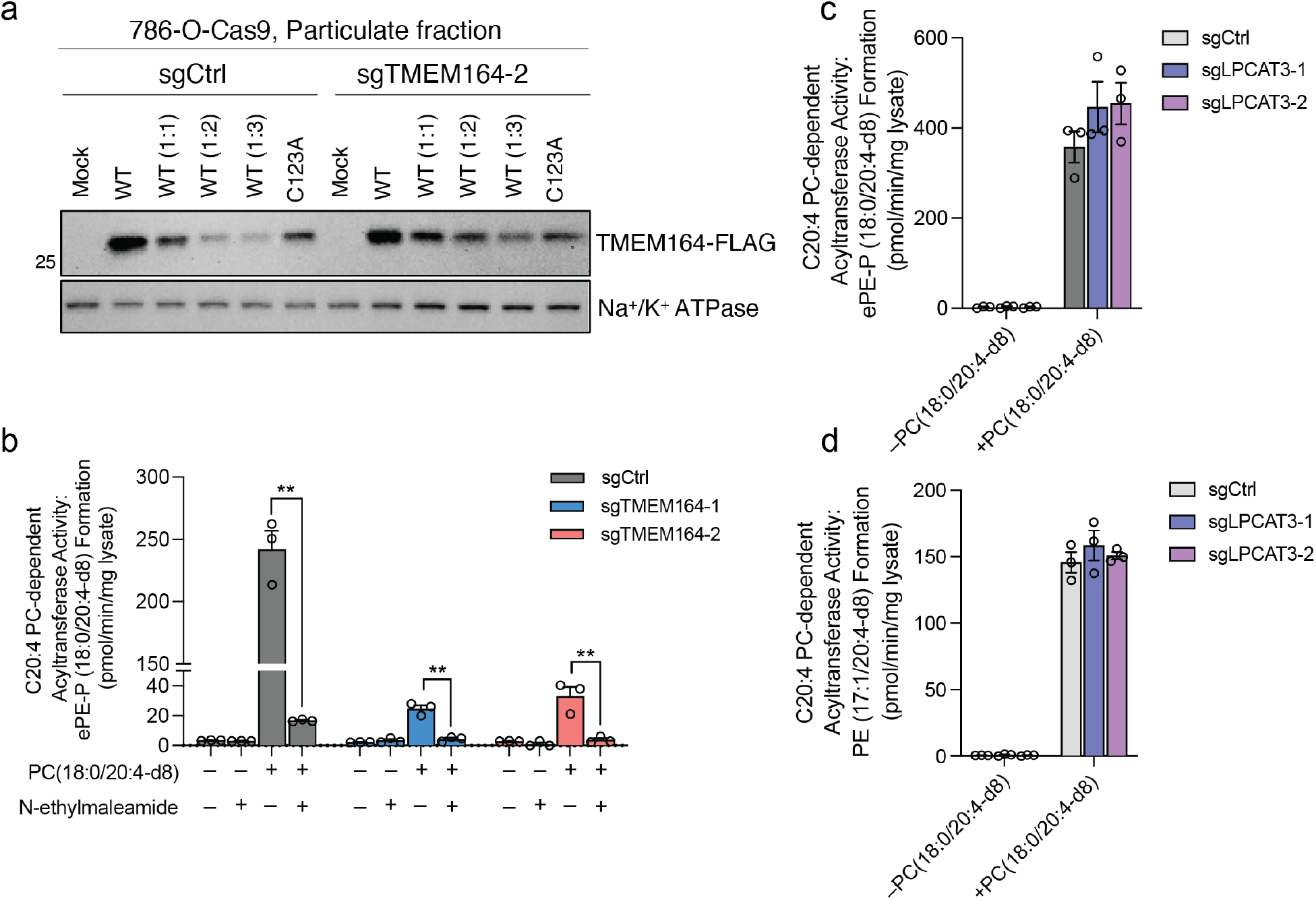
Characterization of C20:4 PC-dependent acyltransferase activity of TMEM164. **a,** Western blot of membrane lysates from sgCtrl and sgTMEM164-2 786-O-Cas9 cells recombinantly expressing C-terminally FLAG-tagged WT-TMEM164 or a C123A-TMEM164 mutant. WT-TMEM164 lysates were serially diluted with mock lysate to identify a dilution (1:1) that matched the expression of C123A-TMEM164. Data are from a single experiment representative of two independent experiments. **b**, Membrane lysates from sgCtrl or sgTMEM164 786-O-Cas9 cells were treated with *N*-ethylmaleimide (NEM; 200 μM, 60 min) prior to exposure to diester PC(18:0/20:4-d8) and lyso-ePE-P(18:0) substrates and measurement of ePE-P(18:0/20:4-d8) product. **c, d**, Measurement of (**c**) ePE-P(18:0/20:4-d8) and (**d**) diester PE(17:1/20:4-d8) formation in membrane lysates from sgCtrl and sgLPCAT3 786-O-Cas9 cells using diester PC(18:0/20:4-d8) and ePE-P(18:0) or LPE(17:1) as donor and acceptor substrate, respectively. For **b-d**, membrane lysates (1 μg/μL) were treated with 50 μM each of donor and acceptor substrates for 1 h at 37° C prior to analysis. For **b-d** data represent mean values ± S.E.M from three independent experiments per group. ** p < 0.001 (Two-sided Student’s t-test performed relative to sgCtrl cells).

**Extended Data Figure 7.**
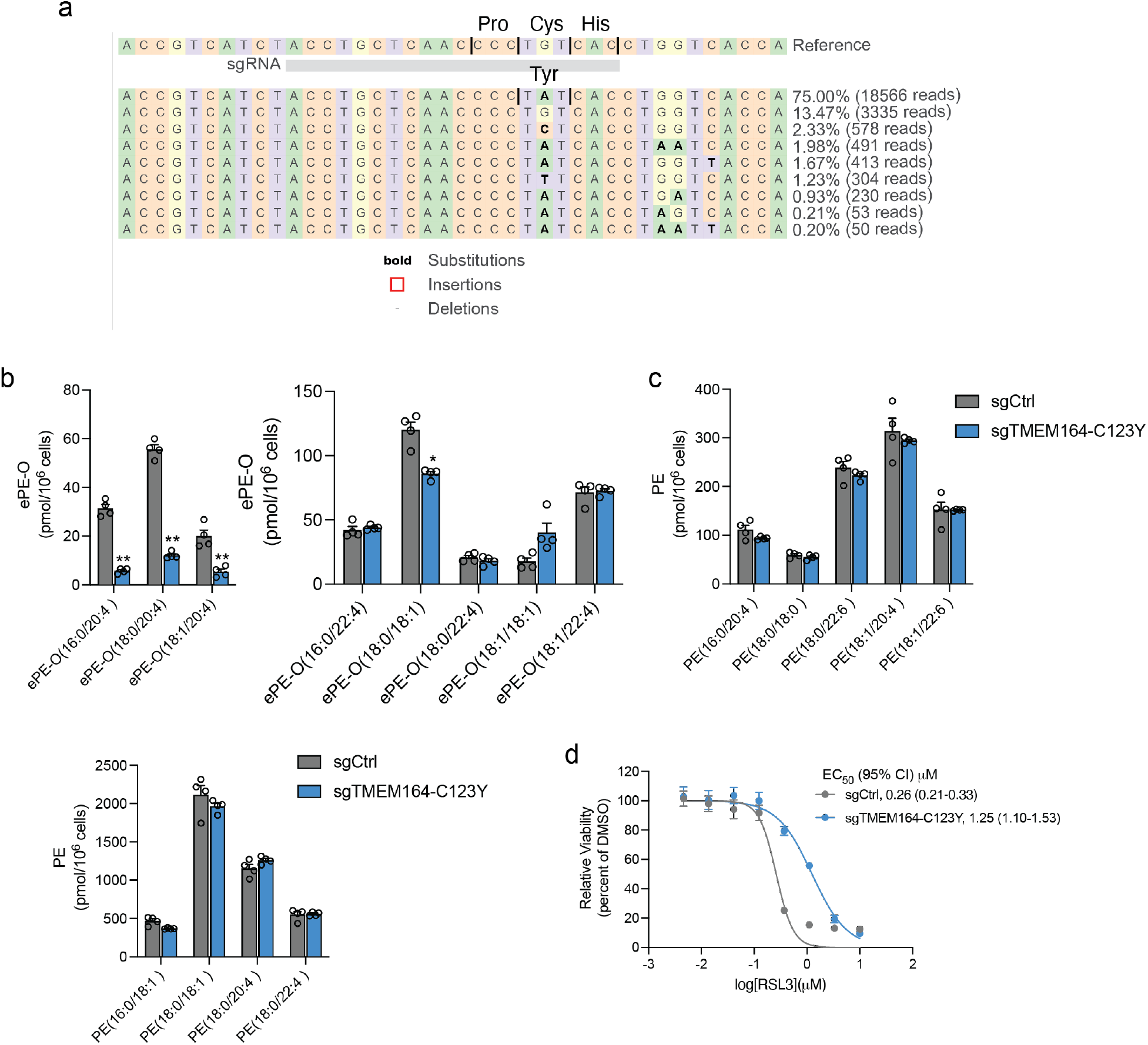
Characterization of sgTMEM164-C123Y base-edited cells. **a**, Confirmation of base-editing of TMEM164 genomic DNA in 786-O cells at the sgRNA target site for sgTMEM164-C123Y base-edited cell population by next-generation sequencing analysis. **b**, Measurement of C20:4 (left) and additional (right) ePE-O lipids in sgCtrl and sgTMEM164-C123Y base-edited cell populations. **c**, Measurement of diester PE lipids in sgCtrl and sgTMEM164-C123Y base-edited 786-O cell populations. **d**, Cell viability of sgCtrl and sgTMEM164-C123Y base-edited cell populations treated with the indicated concentrations of the GPX4 inhibitor RSL3 measured at 24 h post-treatment. For **b** and **c** data represent mean values ± S.E.M from four independent experiments per group. For **d** data represent mean values ± S.E.M. from two independent experiments with two-three technical replicates per experiment. *p<0.01 and ** p < 0.001 (Two-sided Student’s t-test performed relative to sgCtrl cells).

**Extended Data Figure 8.**
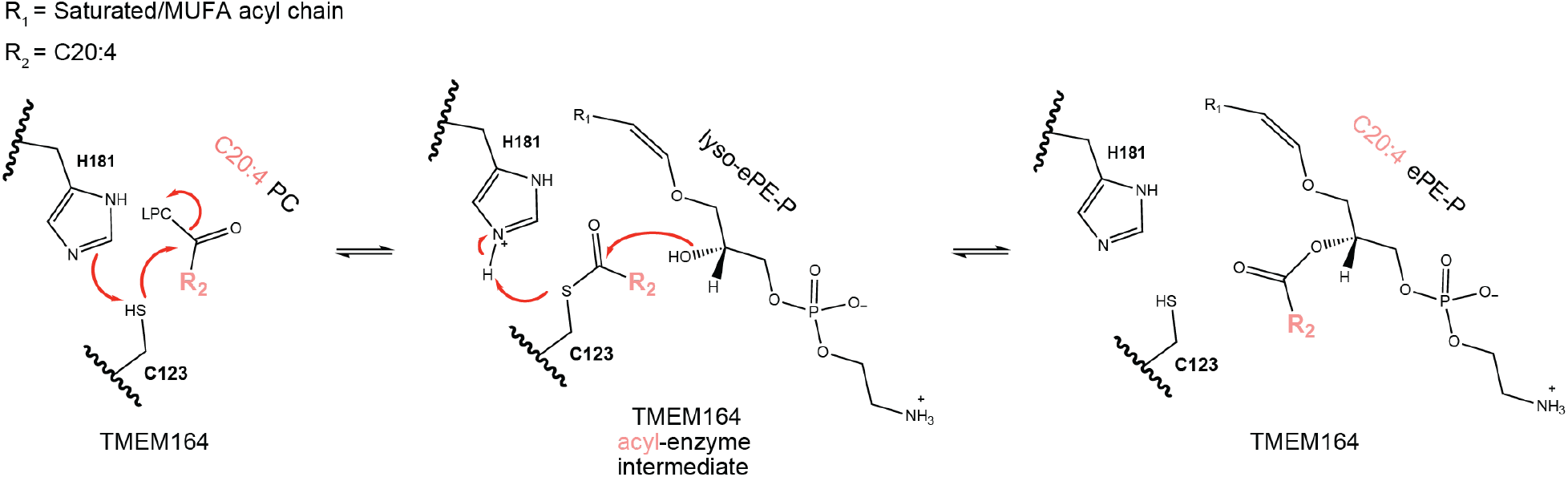
Proposed catalytic mechanism for C20:4 PL-dependent acyltransferase activity of TMEM164. LPC, lysophosphatidylcholine.

**Extended Data Figure 9.**
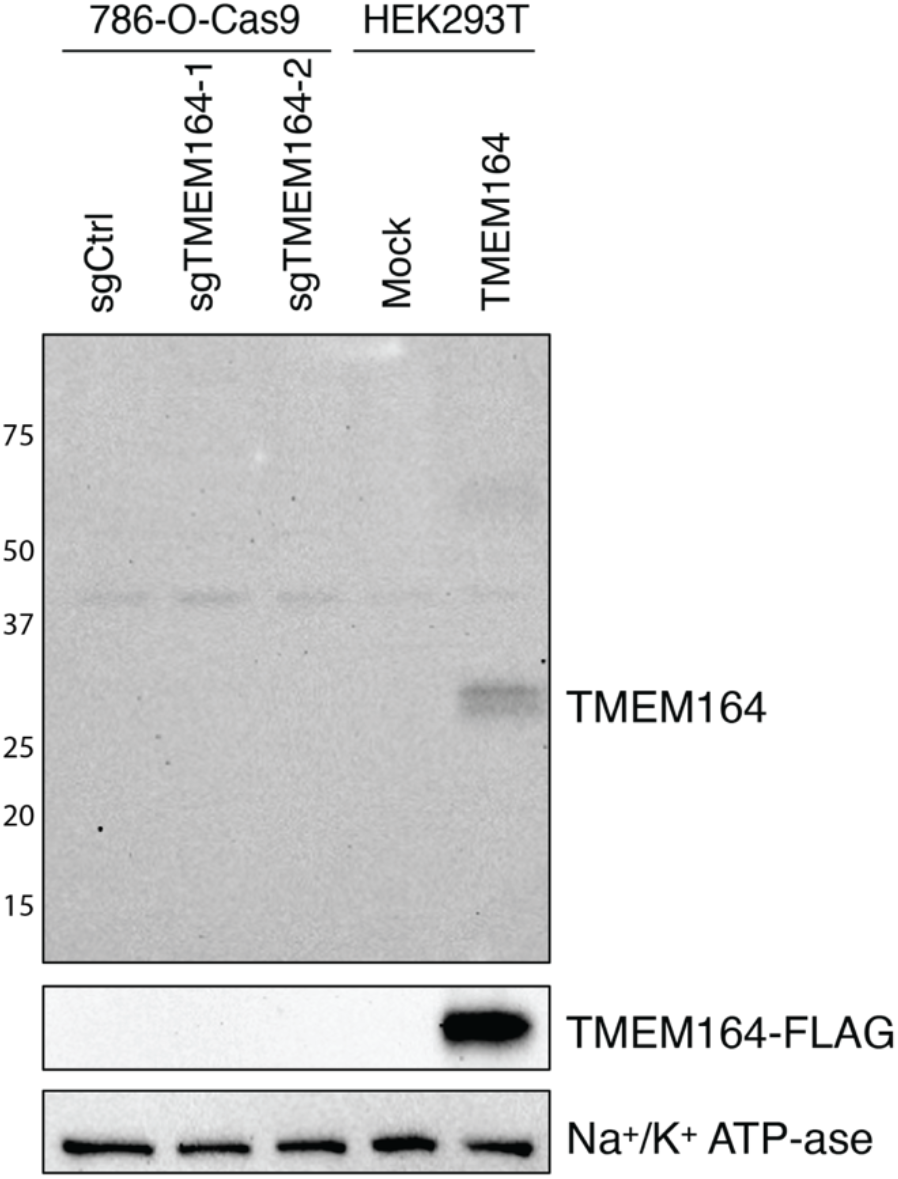
Western blots of membrane lysates from sgCtrl and sgTMEM164 786-O-Cas9 or HEK293T cells expressing mock or TMEM164-FLAG using commercial anti-TMEM164 (upper blot; Thermo Scientific, PA5-58540) or anti-FLAG (middle blot; Abcam, ab2493) antibodies.

