## Supplementary material for "TMEM164 is an acyltransferase that forms ferroptotic polyunsaturated ether phospholipids": Methods

### MATERIALS AND METHODS

#### Materials

All chemicals were obtained from Sigma Aldrich unless indicated otherwise. All lipids were purchased from Avanti Polar lipids or Cayman Chemical Company. RSL3 and ML210 were purchased from SelleckChem.

#### Cell lines

Cell lines were purchased from ATCC and tested negative for mycoplasma contamination. HEK293T cells were maintained at 37°C with 5% $CO_2$ in DMEM (Corning, 15-013-CV) supplemented with 10% (v/v) fetal bovine serum (FBS, Omega Scientific), penicillin (100 U/mL), streptomycin (100 µg/mL) and L-glutamine (2mM). 786-O derived cells were maintained at 37°C with 5% $CO_2$ in RPMI (Corning, 15-040-CV) supplemented with 10% (v/v) FBS, penicillin (100 U/mL), streptomycin (100 µg/mL) and L-glutamine (2 mM).

#### C20:4 PL-dependent acyltransferase LC-MS based substrate assay

HEK293T, 786-O-Cas9 sgCtrl or sgTMEM164 cell lysates were lysed in assay buffer A [30 mM Tris-HCl (pH 7.4), 5 mM EDTA, 0.12 M NaCl] and complete protease inhibitor, EDTA-free (Roche) using a probe sonicator (Branson Sonifier model 250) with 8 pulses (30% duty cycle, output setting = 4) and membrane fraction obtained by centrifugation at 100,000 x g for 45 min at 4°C. Membrane lysates were prepared by sonicating cell pellets in assay buffer A and total protein concentrations were determined using the Bio-Rad DC protein assay kit. Membrane lysates were diluted to 1 mg/mL using assay buffer A. 66 µM of both the lipid acyl acceptor (e.g., C18:0 lyso-ePE-P) and lipid acyl donor (e.g., C18:0/20:4-d8 PC) were prepared by resuspending dried down stocks in assay buffer A. Assay was performed by combining 60 µL of membrane lysates with 190 µL of 66 µM lipid substrate solutions (50 µM final concentration) and incubating at 37°C with shaking (200 rpm) for 60 min. The reaction was quenched by the addition of 450 µL 2:1 $CHCl_3$/MeOH (v/v) with 1 nmol C17:0/20:4 PE as an internal standard. The mixture was vortexed and centrifuged at 2,000 x g for 10 min at 4°C to separate the aqueous and organic phases. The organic phase was extracted and dried down under $N_2$ stream. Lipids were suspended in 100 µL 2:1 $CHCl_3$/MeOH and stored at -80°C until analyzed. The organic phase was analyzed by LC/MS–based multiple reaction monitoring (MRM) (Agilent Technologies 6460 or 6470 Triple Quad). MS analysis was performed using negative mode ESI with the following parameters: drying gas temperature, 350°C; drying gas flow, 9 l/min; nebulizer pressure, 50 Ψ; sheath gas temperature,

375°C; sheath gas flow, 12 l/min; fragmentor voltage, 100 V; and capillary voltage, 3.5 kV. The separation of the analyte was achieved using a 50 mm × 4.6 mm 5 μm Gemini C18 column (Phenomenex) coupled to a guard column (Gemini: C18: 4 × 3 mm). The LC solvents were as follows: buffer A, $H_2O$/MeOH (95:5, v/v) with 0.1% $NH_4OH$ (v/v); and buffer B, iPrOH/MeOH/$H_2O$ (60:35:5, v/v/v) with 0.1% $NH_4OH$ (v/v). The LC gradient following after injection: 20% B at 0.1 mL/min for 5 min; then increase to 85% B at 0.4 mL/min for 15 min; increase to 100% B at 0.5 mL/min for 5 min, run at 100% B at 0.5 mL/min for 6 min; then go back to 20% B and equilibrate at 0.5 mL/min for 3 min. ePE-P(18:0/20:4-d8) (758.5 → 311.3), ePE-P(18:0/20:4-d11) (761.5 → 314.3), ePE-P(18:0/18:0-d35) (765.5 → 318.3), ePE-P(18:0/18:1) (733.5 → 286.3), PE(17:0/20:4) (*m/z* 750.5 → *m/z* 303.3), PE(17:1/20:4-d8) (*m/z* 758.5 → *m/z* 311.3), and PE(17:0/20:4) (*m/z* 752.5 → *m/z* 303.3) were measured.

**Coenzyme A-dependent acyltransferase LC-MS based substrate assay**

HEK293T cell lysates over-expressing hTMEM164-FLAG or hLPCAT-FLAG were lysed using a probe sonicator (Branson Sonifier model 250) with 15 pulses (30% duty cycle, output setting = 3) and membrane fraction was obtained by centrifugation at 16,300 x g for 45 min at 4°C. Membrane lysates were prepared by sonicating cell pellets in assay buffer B [10 mM Tris-HCl (pH 7.4), 1 mM EDTA, 0.15 M NaCl] and total protein concentrations were determined using the Bio-Rad DC protein assay kit. 50 μM each of C17:1 LPE and C20:4-CoA was added to the membrane lysates diluted in assay buffer B (0.01 mg/mL) and allowed to incubate for 10 min at room temperature. The reaction was quenched by the addition of 300 μL 2:1 $CHCl_3$/MeOH (v/v) with 1 nmol C12:0/12:0 PE as an internal standard. The mixture was vortexed and centrifuged at 2,000 x g for 5 min at 4°C to separate the aqueous and organic phase. The organic phase was analyzed by LC/MS–based multiple reaction monitoring (MRM) (Agilent Technologies 6460 or 6470 Triple Quad). MS analysis was performed using ESI with the following parameters: drying gas temperature, 350°C; drying gas flow, 9 l/min; nebulizer pressure, 45 Ψ; sheath gas temperature, 375°C; sheath gas flow, 12 l/min; fragmentor voltage, 100 V; and capillary voltage, 3.5 kV. The separation of the analyte was achieved using a 50 mm × 4.6 mm 5 μm Gemini C18 column (Phenomenex) coupled to a guard column (Gemini: C18: 4 × 3 mm). The LC solvents were as follows: buffer A, $H_2O$/MeOH (95:5, v/v) with 0.1% $NH_4OH$ (v/v); and buffer B, iPrOH/MeOH/$H_2O$ (60:35:5, v/v/v) with 0.1% $NH_4OH$ (v/v). The LC gradient following after injection: 20% B at 0.8 mL/min for 6 min; then increase to 100% B at 1 mL/min for 2 min; decrease to 20% B at 1 mL/min for 1 min; then equilibrate at 20% B at 0.8 mL/min for 0.5 min. C17:1/20:4 PE (*m/z*

750.5 → *m/z* 303.3) and C12:0/12:0 PE (internal standard, *m/z* 578.3 → *m/z* 199.2) were measured.

**Metabolomic analysis for cell samples**

Cells were washed twice with cold DPBS, and the total cell metabolome was extracted in 4 mL 2:1:1 $CHCl_3$/MeOH/DPBS (v/v/v) solution containing the internal standard mix (100 pmol C12:0/12:0 PC and 100 pmol C12:0/12:0 PE). The mixture was vortexed vigorously and centrifuged at 2,000 x g for 5 min at 4°C. The bottom organic phase was collected, and the remaining aqueous phase was acidified with 100 µL formic acid and re-extracted by the addition of 2 mL $CHCl_3$. Both organic extracts were pooled, dried down under $N_2$ stream, and reconstituted in 150 µL 2:1 $CHCl_3$/MeOH (v/v) for LC/MS analysis.

Metabolites analyzed in this study were quantified using LC/MS–based multiple reaction monitoring (MRM) methods (Agilent Technologies 6460 or 6470 Triple Quad). MS analysis was performed using ESI with the following parameters: drying gas temperature, 350 °C; drying gas flow, 9 l/min; nebulizer pressure, 45 Ψ; sheath gas temperature, 375 °C; sheath gas flow, 12 l/min; fragmentor voltage, 100 V; and capillary voltage, 3.5 kV. The MRM transitions for the targeted LC/MS analysis are presented in the compiled lipidomics data spreadsheet. The separation of metabolites was achieved using a 50 mm × 4.6 mm 5 µm Gemini C18 column (Phenomenex) coupled to a guard column (Gemini: C18: 4 × 3 mm). For negative mode analysis, $H_2O$:MeOH (95:5, v/v) with 0.1% $NH_4OH$ (v/v) and iPrOH:MeOH:$H_2O$ (60:35:5, v/v) with 0.1% $NH_4OH$ (v/v) were used as buffer A and B, respectively. For positive mode analysis, 20 mM ammonium acetate in $H_2O$ and 20 mM ammonium acetate in MeOH were used as buffer A and B, respectively. The LC gradient for negative mode analysis was the following after injection: 20% B at 0.1 mL/min for 5 min; then increase to 85% B at 0.4 mL/min for 15 min; increase to 100% B at 0.5 mL/min for 5 min, run at 100% B at 0.5 mL/min for 2 min; then go back to 20% B and equilibrate at 0.5 mL/min for 5 min. The LC gradient for positive mode analysis was the following after injection: start from 75% B and increase to 99% B at 0.35 mL/min for 22 min; run at 99% B for 17 min; then go back to 75% B and equilibrate for 3 min. Lipid species were quantified by measuring areas under the curve in comparison to the corresponding internal standards and then normalizing to the cell numbers.

Isobaric ePE-O and ePE-P lipids were separated by ~1.5 min retention time difference, with ePE-O eluting before ePE-P as previously reported[1,2]. Treatment of lipid extracts with 10%

(v/v) of formic acid or HCl (3N) in a pilot experiment was used to confirm the difference in retention time between ePE-O and ePE-P species (**Extended Data Fig. 3**). ePC-O and ePC-P species were not distinguished in this study, as has been described in previous protocols[3].

**Generation of CRISPR-mediated TMEM164- and LPCAT3-null 786-O cells**

sgTMEM164 and sgLPCAT3 786-O-Cas9 cells were generated as described previously. Briefly, 786-O cells were transduced with pLX-311-Cas9 vector (Addgene 96924), which results in constitutive expression of SpCas9 and contains the blasticidin S-resistance gene. Lentiviral sgRNAs targeting human TMEM164 and LPCAT3 along with a control sgRNA were cloned into the pXPR_BRD050 constitutive sgRNA expression vector. The sequences for the sgRNAs are described below (sequences are 5' to 3').

sgCTRL-fwd:caccgGCGAGGTATTCGGCTCCGCG
sgCTRL-rev:aaacCGCGGAGCCGAATACCTCG

sgTMEM164-1-fwd: caccgCATCCTGAGGCAGACGAAGG
sgTMEM164-1-rev: aaacCCTTCGTCTGCCTCAGGATGC

sgTMEM164-2-fwd:caccGCCTGTGCCTGACCTTCGGGG
sgTMEM164-2-rev:aaacCCCCGAAGGTCAGGCACAGGc

sgLPCAT3-1-fwd: caccGAGACTCAGGCGCTTGAGAGC
sgLPCAT3-1-rev: aaacGCTCTCAAGCGCCTGAGTCTC

sgLPCAT3-2-fwd: caccGACTGAAGCACAATACACAGC
sgLPCAT3-2-rev: aaacGCTGTGTATTGTGCTTCAGTC

sgRNA-encoding plasmids were co-transfected with dr8.9 envelope and VSV-G packaging plasmids into 500,000 HEK293T cells in 6cm dishes using the Fugene6 transfection reagent (Promega). Virus-containing supernatants were collected 48 h after transfection and used to infect target 786-O-Cas9 cells in the presence of 10 mg/mL polybrene (Santa Cruz). The cells were allowed to recover for 48 h. Puromycin (2 μg/mL) and Blasticidin (10 μg/mL) was then added to the cells, and the cells were expanded. Genetic alterations in sgLPCAT3 cells were previously reported[4], while genetic alterations in sgTMEM164 cells were confirmed by next-generation sequencing analysis as described below.

**Generation of sgTMEM164-C123Y base-edited 786-O cells**

Plasmid containing the cytidine deaminase conjugated to dCas9 was a gift from the laboratory of David Liu. sgRNA targeting C123 on TMEM164 as well as a control sgRNA targeting GFP was cloned into the vector. The sequences are described below (sequences are 5' to 3').

sgCTRL-fwd: caccgGCGAGGTATTCGGCTCCGCG
sgCTRL-rev: aaacCGCGGAGCCGAATACCTCG

sgTMEM164-C123Y-fwd: caccgGTGACAGGGGTTGAGCAGGT
sgTMEM164-C123Y-rev: aaacACCTGCTCAACCCCTGTCACc

sgRNA-encoding plasmids were co-transfected with dr8.9 envelope and VSV-G packaging plasmids into 500,000 HEK293T cells in 6cm dishes using the Fugene6 transfection reagent (Promega). Virus-containing supernatants were collected 48 h after transfection and used to infect target 786-O cells in the presence of 10 mg/mL polybrene (Santa Cruz). The cells were allowed to recover for 48 h. Puromycin (2 µg/mL) was then added to the cells, and the cells were expanded. Genetic alterations in sgTMEM164-C123Y cells were confirmed by next-generation sequencing analysis as described below.

**Targeted genomic sequencing and data analysis**

Genomic sites of interested were amplified and indexed using a two-step PCR method as previously described[5]. Briefly, cells were lysed using 10 mM Tris pH 7.5, 0.5% Tween 20, 0.02% SDS plus 20 µg/ml freshly added proteinase K and were incubated at 55°C for 2 hr before heat inactivation at 90°C for 30 min. Primers containing homology to targeted region and illumina adapters were used for PCR1 (sequences below, 5' to 3'). Specifically, in each reaction, 5 µl of genomic DNA extract, 12.5 µl Phusion PCR master mix, 1.25 µl of 10 µM forward and 1.25 µl of 10 µM reverse primers were added to a total volume of 25 µl. The PCR1 reactions were carried out as follows: 98°C for 3 min, then 30 cycles of (20s of 98°C, 25s of 60°C, 25s of 72°C), followed by final extension at 72°C for 2 min. PCR1 products were cleaned using Ampure beads (Beckman) according to manufacturer's instructions and were eluted in 20 µl of water. For each PCR2 reaction, 5 µl of PCR1 product, 12.5 µl Phusion PCR master mix, 1.25 µl of 10 µM forward and 1.25 µl of 10 µM reverse index primers were added to a total volume of 25 µl. The PCR2 reactions were carried out as follows: 98°C for 3 min, then 12 cycles of (20s of 98°C, 20s of 60°C, 25s of 72°C), followed by final extension at 72°C for 2 min. The PCR2 products were then pooled and cleaned using Ampure beads. The library was quantified using PicoGreen dsDNA assay kits (ThermoFisher) and sequenced on an Illumina Miniseq instrument with 30% PhiX spike-in. Samples reads were demultiplexed based on combinatorial dual indexes. The genome editing quantification was performed using CRISPResso2 (https://crispresso.pinellolab.partners.org)[6].

Forward Sequencing Primer: ACACGACGCTCTTCCGATCTGGAGAGCCTGAGCAAGAATC
Reverse Sequencing Primer: CTTGGCACCCGAGAATTCCAGGATGAGGTGCTGTACACTATG

**Western blot analysis**

Cell pellets were lysed in DPBS and complete protease inhibitor, EDTA-free using a probe sonicator (Branson Sonifier model 250) with 15 pulses (30% duty cycle, output setting = 4). Proteins were resolved by SDS–PAGE, transferred to 0.45 µM nitrocellulose membranes (Thermo Fisher) which were blocked with 5% milk in TBST buffer (20 mM Tris-HCl 7.6, 150 mM NaCl with 0.1% tween 20). Primary antibodies were used at the following concentrations: 1:2,000 anti-$Na^+/K^+$-ATPase (Cell Signaling, no. 3010), 1:2,000 anti-FLAG-HRP conjugated (Abcam, ab2493), 1:2,000 anti-TMEM164 polyclonal (Thermo Scientific, PA5-58540). Blots were incubated with primary antibodies in 5% milk in TBST at 4°C overnight. Following another TBST wash (5 times, 5 min), membranes were incubated with 1:5,000 anti-rabbit IgG, HRP conjugated secondary antibody in 5% milk in TBST (Cell Signaling, no. 7074S) at room temperature for 1 h. Membranes were washed with TBST (5 times, 5 min) and developed with ECL western blotting detection reagent kit (Thermo Scientific) and detected with Bio-Rad ChemiDoc MP imaging system.

**Recombinant expression of TMEM164 wild type and C123A mutant in sgCtrl and sgTMEM164-2 786-O-Cas9 cells**

Full length C-terminally FLAG-tagged TMEM164 and C123A mutant were obtained by ordering a gBlock gene fragment from Integrated DNA Technologies with a silent point mutation at the PAM site for sgTMEM164-2 and attB1/attB2 sites flanking the gene. Gene fragments were cloned into the gateway compatible pLV416 vector by standard gateway cloning. TMEM164-encoding plasmids were co-transfected with dr8.9 envelope and VSV-G packaging plasmids into 4 million HEK293T cells in 10cm dishes using the Fugene6 transfection reagent (Promega). Virus-containing supernatants were collected 48 h after transfection and used to infect sgCtrl or sgTMEM164-2 786-O-Cas9 cells in the presence of 10 mg/mL polybrene (Santa Cruz). The cells were allowed to recover for 48 h. Geneticin (0.8 mg/mL) was then added to the cells, and the cells were expanded.

**Recombinant expression of TMEM164 wild type and C123 mutants by transfection of HEK293T cells**

TMEM164 C123X mutants were obtained by ordering gBlock gene fragments from Integrated DNA Technologies with attB1/attB2 sites flanking the gene. Gene fragments were cloned into the gateway compatible pLV416 vector using standard gateway cloning techniques. HEK293T cells were transiently transfected with TMEM164 WT or C123X mutants using PEI and cells were collected for analysis 48 h later.

***In situ* metabolic labeling assay**

786-O-Cas9 sgCtrl and sgTMEM164-2 cells were plated at 500,000 cells in a 6cm dish the day prior to the assay and then treated with DMSO, C20:4-d8 FFA or C18:1-d9 FFA (Cayman) (25 μM) for 4 h. Following treatment, cells were collected, washed twice with ice-cold PBS and pellets were stored at -80°C for further analysis. Metabolomics analysis was performed in negative mode as described above.

**Ferroptosis assay**

786-O cells were plated in white, clear-bottom 96-well plates (Greiner) at 5,000 cells per well the day prior to the assay. On the day of the assay, DMSO or a serial dilution of RSL3 or ML210 (3-fold dilution starting from 10 μM, 8 concentrations) were added to the wells and allowed to incubate for 24 h. At 24 h, the plates were removed from the incubator, 50 μL of CellTiter-GLO (CTG) (Promega) was added to each well and allowed to incubate for 30 min at room ttemperature. Luminescence was measured on a Clariostar plate reader (BMG Labtech).

**Computational and structural analyses.**

The initial discovery of a greater superfamily beyond AIG1/ADTRP—previously identified as novel FAHFA hydrolases in lipid metabolism—was done with sensitive sequence search algorithms, iterative PsiBLAST and HHPRED, using servers located in the BLAST suite at the NCBI (http://blast.ncbi.nlm.nih.gov) and as part of the bioinformatics Toolkit at the MPI-Tübingen (http://toolkit.tuebingen.mpg.de/tools/hhpred), respectively. In particular, HHPRED considers PsiPRED-generated secondary structure predictions in addition to the iteratively-generated PSSM sequence profiles, and these enhanced search tools can be aimed at diverse proteome-level collections of sequences, as well as various structure and domain databases, to detect distant matches. In this manner with HHPRED, the human AIG1 sequence locates AIG1/ADTRP transmembrane enzyme homologs in *D.melanogaster* and *S.cerevisiae* (consistent with the PFAM PF04750 family cluster; PFAM databases are maintained at http://pfam.xfam.org), and also finds more distant, partial matches with prokaryotic YwaF members (like *B.subtili*s and *C.difficile*, members of PF09529 family) that show alignment of predicted TM helices and overlap of potential catalytic dyads—with an invariant, C-terminal His, and a variable (Thr, Cys or Ser), more N-terminal, nucleophile. Iterative PsiBLAST runs that reach into sequence families absent from HHPRED, further link the YwaF proteins to members of the Hypothetical clan (principally in archaea, also featuring variable nucleophile catalytic dyads) and the more distant YpjA clan (PF7187, prinicipally Ser/His catalytic dyads). These latter enzymes were used to query eukaryotic proteomes with HHPRED, and locate the orphanTMEM164 sequence in humans (as well as *D.melanogaster* and *C.elegans*), building a distant homology link to PFAM family PF14808 that exclusively use a Cys/His catalytic dyad.

The structural implications of these weak sequence similarity HHPRED and PsiBLAST matches was tested with the best 2$^{nd}$ generation structure prediction program, TrRosetta, using the server at Nankai University (http://yanglab.nankai.edu.cn/trRosetta), that generated 3D structure models of AIG1/ADTRP, YwaF and YpjA, Hypothetical and TMEM164 proteins, and asserted that they had a common 6TM core fold (**Fig. 4a,b**, **Extended Data Fig. 5a,b**) decorated by N- and C-terminal extensions of variable length. The Thr/His catalytic dyad in human AIG1 was distributed across TM2 and TM4, respectively, while the Cys/His dyad in human TMEM164 likewise mapped to structurally equivalent locations in TM2 and TM4. The advent of the new deep-learning-based AlphaFold2 algorithm (http://www.deepmind.com/research/highlighted-research/alphafold), allowed much more precise modeling of representative members of the 5 linked families of sequences with the versatile and fast ColabFold implementation (http://github.com/sokrypton/ColabFold), that reasserted even more accurately, the presence of a common 6TM core fold bearing the enzymatic machinery of this emerging membrane protein superfamily (**Fig. 4c**, **Extended Data Fig. 5e**). In many cases now, AlphaFold2 models of superfamily members can be directly retrieved from the AlphaFold database maintained at the

EBI (https://alphafold.ebi.ac.uk), that is now incorporated into most UniProt files (https://www.uniprot.org).

Fast searches of the extensive AlphaFold model databases—organized by proteomes—using Rupee and FoldSeek servers (respectively at http://ayoubresearch.com and https://search.foldseek.com) validated the homology links of the 5 sequence families by showing significant fold relationships. In addition, these fold-level screens discovered an additional integral membrane enzyme family headlighted by S.cerevisiae and A.thaliana GPC1 enzymes (members of PF10998 family) at very low, ~10% sequence identity, further sharing the same Cys/His catalytic dyad of the human TMEM164 chain, that had eluded detection by the HHPRED searches. AlphaFold2 models of GPC1 enzymes revealed the same core 6TM fold with catalytic residues in TM2 and TM4, with the largest N- and C-terminal extensions (**Fig. 4b, Extended Data Fig. 5d**).

The low level of sequence identity between the now six branches of novel enzymes in the AIG and ADTRP/TMEM164/GPC1/YwaF and YpjA (or ATGY) superfamily was insufficient to gauge their evolutionary relationship, so the slow but rigorous DALI algorithm (http://ekhidna2.biocenter.helsinki.fi) was used to perform an all-against-all matching of the AlphaFold2-generated models. The average linkage clustering of DALI Z-scores (**Extended Data Fig. 5c**), a measure of structural similarity, was used to generate dendrograms as conventional linear trees (**Extended Data Fig. 5b**) or hyperbolic trees (**Fig. 4c**, made with the HyperTree program from http://kinase.com/tools/HyperTree.html) of the ATGY superfamily, showing clear separation of the six branches, with their particular and divergent catalytic dyad preferences.

Closer structural analysis of the TMEM164 fold was performed with CavityPlus (http://www.pkumdl.cn/cavityplus) and CASTp programs (http://sts.bioe.uic.edu), to delineate the size and shape of the internal cavity in the 6TM helical core (**Fig. 4d**, **Extended Data Fig. 5e**). CavityPlus also links to CavPharmer that additionally builds a pharmacophore ‘skeleton’ (by analysis of residue constellations surrounding the cavity) that fits within the cavity and abuts the catalytic dyad and is highly suggestive of a phospholipid ligand (which could be a substrate or product).

### $EC_{50}$ calculation

For the acyltransferase assays, acyltransferase activities were determined as a ratio of the area under the peak of the expected product over the area under the peak of either C17:0/20:4 PE (internal standard, *m/z* 752.5 → *m/z* 303.3) or C12:0/12:0 PE (internal standard, *m/z* 578.3 → *m/z* 199.2). Data were then normalized by dividing the acyltransferase activity by the amount of membrane proteome used and the length of the assay. For the ferroptosis assay, relative cell viability was determined by dividing the CTG value from RSL3/ML210 treated wells by the CTG value from the average of the DMSO treated wells and multiplying by 100. $EC_{50}$ values were determined by plotting a log(inhibitor) vs. normalized response, and the dose-response curves were generated using the Prism software (GraphPad Software, Inc.).

**Statistics**

Statistical analyses were performed using GraphPad Prism (GraphPad Software, Inc.). All data are shown as mean values ± SEM or SD and two-sided Student's t-test was used to perform statistical analyses. A p-value of < 0.01 was considered statistically significant for this study.

**Supporting Information**.

The following files are available free of charge. Supplementary figures and tables and Materials and methods (PDF). Compiled lipidomics data (excel).
